# Evolutionary and multi-omic divergences associated with the extreme sexual size dimorphism in spiders

**DOI:** 10.64898/2026.09.17.752500

**Authors:** Zheng Fan, Lu-Yu Wang, Jia-Xin Gao, Tian-Yu Ren, Wen-Hui Wu, Wei Pu, Jin-Xia Kong, Yong-Gang Hu, Zhi-Sheng Zhang, Chao Tong

## Abstract

Extreme sexual size dimorphism (SSD) represents one of the most striking forms of phenotypic divergence between the sexes, yet the genomic basis of extreme body-size dimorphism remains poorly understood. Orb-weaving spiders exhibit some of the most pronounced female-biased SSD among terrestrial arthropods, providing an exceptional system for investigating the evolutionary and genomic changes associated with extreme SSD. Here, we combined comparative genomics across 51 spider species with sex-resolved transcriptomic and chromatin profiling in the highly dimorphic Joro spider, *Trichonephila clavata*. Across spiders, we identified SSD-associated evolutionary-rate shifts in genes involved in chromatin regulation, growth, and cuticle structural organization. We further found a significant positive association between JHAMT copy number and SSD, with larger gene repertoires occurring in more extremely dimorphic lineages. In *T. clavata*, expanded JHAMT paralogs showed sex- and tissue-biased expression together with sex-specific differences in chromatin accessibility and local chromatin interactions. Integrated multi-omic analyses further revealed coordinated sex-specific differences in transcription, chromatin accessibility, and chromatin interactions across genes involved in ecdysteroid signaling, insulin and growth regulation, and epidermal and cuticular processes. Altogether, these findings provide insight into the evolutionary and regulatory changes associated with extreme female-biased SSD in spiders.

## Introduction

Sexual size dimorphism (SSD), the difference in body size between females and males, is widespread across animals and reflects sex-specific growth and developmental trajectories. SSD has evolved repeatedly and varies markedly in both direction and magnitude across lineages, ranging from male-biased to extremely female-biased patterns (Fairbairn, 1997; Stillwell et al., 2010; Ceballos et al., 2013; Isaac, 2005; Schoenjahn et al., 2020; Kuntner and Coddington, 2020; Beato et al., 2026).

Arthropods exhibit striking variation in SSD across diverse lineages. Pronounced female-biased SSD has been documented in damselflies such as Ischnura elegans (Abbott and Gosden, 2009; Dudaniec et al., 2022), the blacklegged tick Ixodes scapularis (Ronai et al., 2025), horseshoe crabs such as Limulus polyphemus (Smith et al., 2009; Smith and Brockmann, 2014), and orchid mantises such as Hymenopus coronatus (Svenson et al., 2016). Spiders include some of the most extreme examples of female-biased SSD among terrestrial arthropods, particularly in orb-weaving lineages such as *Trichonephila* and *Nephila* (Hormiga et al., 2000; Kuntner and Coddington, 2020). Previous studies have examined the roles of fecundity, mate searching, and sex-specific life histories in the evolution of spider SSD (Kuntner and Elgar, 2014; Kuntner and Coddington, 2020). Recent work in the extremely size-dimorphic spider *Nephilingis cruentata* further showed that larger female size increases per-clutch fecundity but does not necessarily translate into greater lifetime fecundity, highlighting the complexity of selective explanations for extreme SSD (Kralj-Fišer et al., 2026). Despite expanding genomic resources and recent efforts to characterize sex-chromosome evolution in highly dimorphic spiders (Recknagel et al., 2026), the genomic signatures associated with extreme SSD remain poorly characterized.

Endocrine and growth-regulatory pathways are emerging as important candidates for explaining body-size divergence in arthropods. In the orchid mantis *H. coronatus*, female-biased expression of genes involved in 20-hydroxyecdysone (20E) biosynthesis and Hippo signaling has been associated with divergent body size (Huang et al., 2023). In the wolf spider *Pardosa pseudoannulata*, multiple juvenile hormone acid O-methyltransferase (JHAMT) genes were identified, and knockdown of selected copies increased juvenile body weight and accelerated molting (Yang et al., 2021a). More recent work identified methyl farnesoate, a juvenile-hormone-related compound, and demonstrated effects on molting, further supporting the relevance of endocrine regulation to spider growth while also highlighting differences from canonical insect juvenile-hormone biology (Yang et al., 2022). Together, these studies establish links between endocrine regulation, growth, and sexual differentiation. However, broad comparative analyses connecting endocrine and growth-related genomic variation with the evolution of extreme SSD remain limited, particularly in spiders. It therefore remains unclear how sequence evolution, gene-repertoire changes, and sex-dependent regulation are jointly associated with increasing SSD across spider lineages.

Advances in chromatin profiling and three-dimensional genomics provide additional opportunities to investigate the regulatory basis of phenotypic divergence. Chromatin accessibility identifies potential regulatory regions, whereas chromatin interactions capture aspects of the spatial organization of genes and distal elements (Buenrostro et al., 2013; Bonev and Cavalli, 2016). In *Drosophila melanogaster* and *D. simulans*, sex-biased gene expression is associated with sex-dependent histone-modification patterns, implicating chromatin state in the maintenance of transcriptional differences between the sexes (Nanni et al., 2023). Developmental profiling in *D. pseudoobscura* further revealed associations among chromatin-state transitions, enhancer activity, and three-dimensional genome organization (Ali et al., 2024). Integrating transcriptomic, chromatin-accessibility, and chromatin-interaction data can therefore identify genes showing coordinated sex-specific differences across regulatory dimensions. Whether such regulatory divergence is associated with the evolution of extreme SSD remains largely unexplored.

Here, we sought to identify evolutionary and molecular signatures associated with extreme female-biased SSD in spiders. We combined comparative genomic analyses across 51 spider species with sex-resolved multi-omic profiling of the highly dimorphic Joro spider, *Trichonephila clavata*. Across spiders, we tested whether protein evolutionary rates and gene-family size were associated with variation in SSD. In *T. clavata*, we examined relationships between evolutionary rates and sex-biased tissue expression, and integrated RNA-seq, ATAC-seq, and Hi-C to characterize transcriptional and chromatin differences between adult females and males. Our analyses identify SSD-associated evolutionary-rate shifts in growth-, chromatin-, and structural-related genes, an expansion of the JHAMT repertoire associated with increasing SSD, and coordinated sex-specific transcriptional and chromatin differences in endocrine, growth-related, epidermal, and cuticular genes. Together, these results provide a comparative and regulatory framework for understanding the genomic changes associated with extreme female-biased SSD in spiders.

## Results

### Female-biased SSD is particularly extreme in orb-weaving spiders

We first placed spider sexual size dimorphism (SSD) in a broader arthropod context by comparing adult female and male body sizes across 23 representative species and calculating female-to-male body-size ratios (Figure 1; Table S1). SSD varied markedly among taxa, with the most extreme female-biased values occurring in spiders. Among the species examined, the Joro spider, *Trichonephila clavata*, exhibited the highest SSD ratio (4.52), whereas the orchid mantis, *Hymenopus coronatus*, represented another prominent example of extreme female-biased SSD.

**Figure 1.**
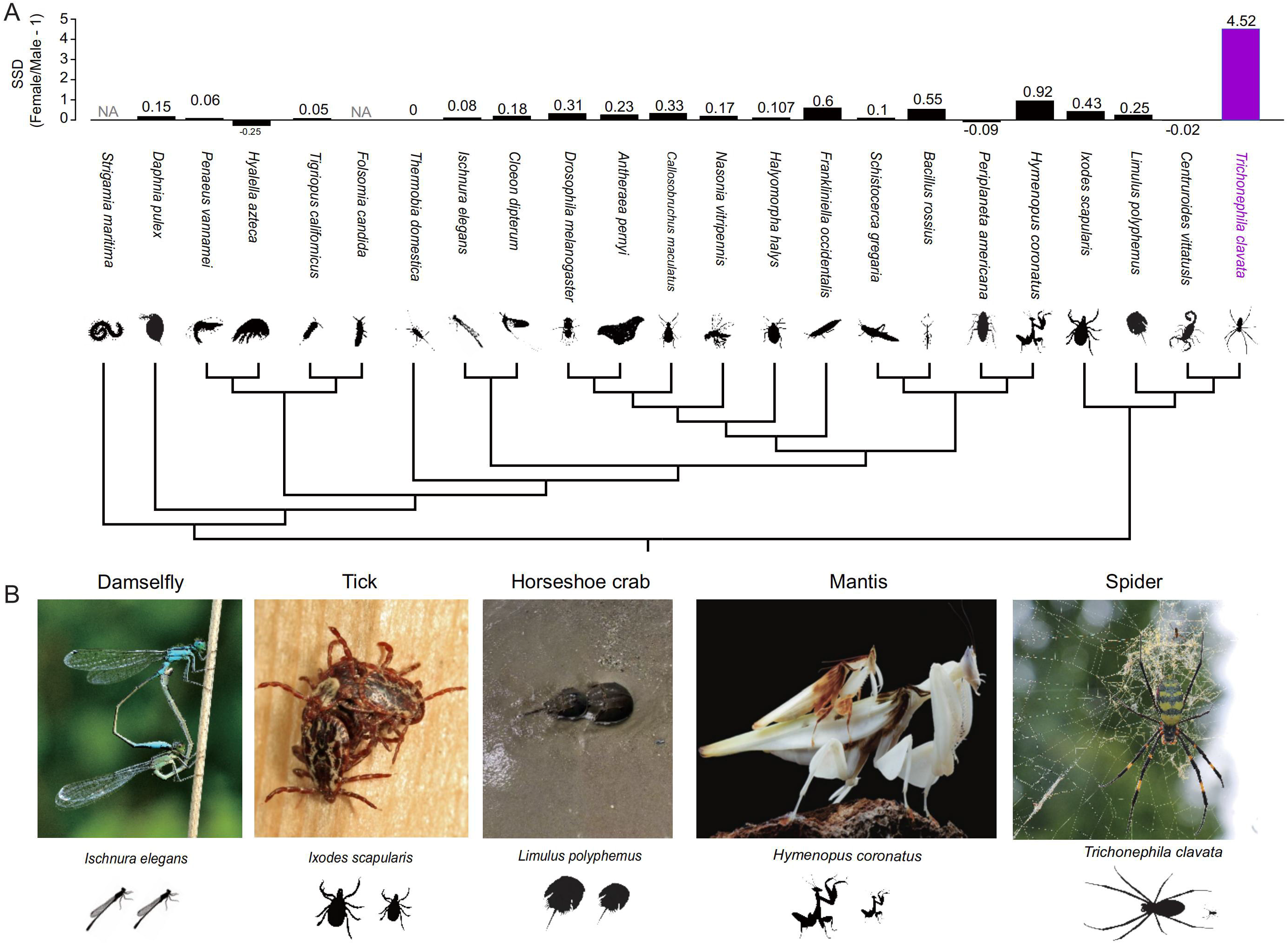
Sexual size dimorphism across representative arthropods. (A) Sexual size dimorphism (SSD) across 23 representative arthropod species shown in a phylogenetic context. SSD was calculated as female body length divided by male body length minus one, such that positive values indicate female-biased SSD and negative values indicate male-biased SSD. *Trichonephila clavata* is highlighted in purple. NA indicates species for which comparable adult female and male body-length measurements were unavailable. (B) Representative examples illustrating female–male size differences in the damselfly *Ischnura elegans*, blacklegged tick *Ixodes scapularis*, horseshoe crab *Limulus polyphemus*, orchid mantis *Hymenopus coronatus*, and Joro spider *T. clavata*. Female and male silhouettes are shown for visual comparison. Photograph credits: *I. elegans*, photograph by Quartl, via Wikimedia Commons (CC BY-SA 3.0); *I. scapularis*, reproduced from Ronai et al. (2025); *L. polyphemus*, photograph by Chao Tong; *H. coronatus*, photograph by Jason Zhu, reproduced from Svenson et al. (2016); *T. clavata*, photograph by Lu-Yu Wang.

We next examined SSD variation across 51 spider species spanning the spider phylogeny (Figure 2A, 2B; Table S2). Female-biased SSD was widespread but differed substantially in magnitude among lineages. The highest values were concentrated primarily within orb-weaving spiders, particularly in nephilines such as *Trichonephila* and *Nephila*. These comparisons establish orb-weaving spiders as a system exhibiting exceptionally pronounced female-biased SSD and provide a phylogenetic framework for identifying genomic changes associated with increasing SSD across spiders.

**Figure 2.**
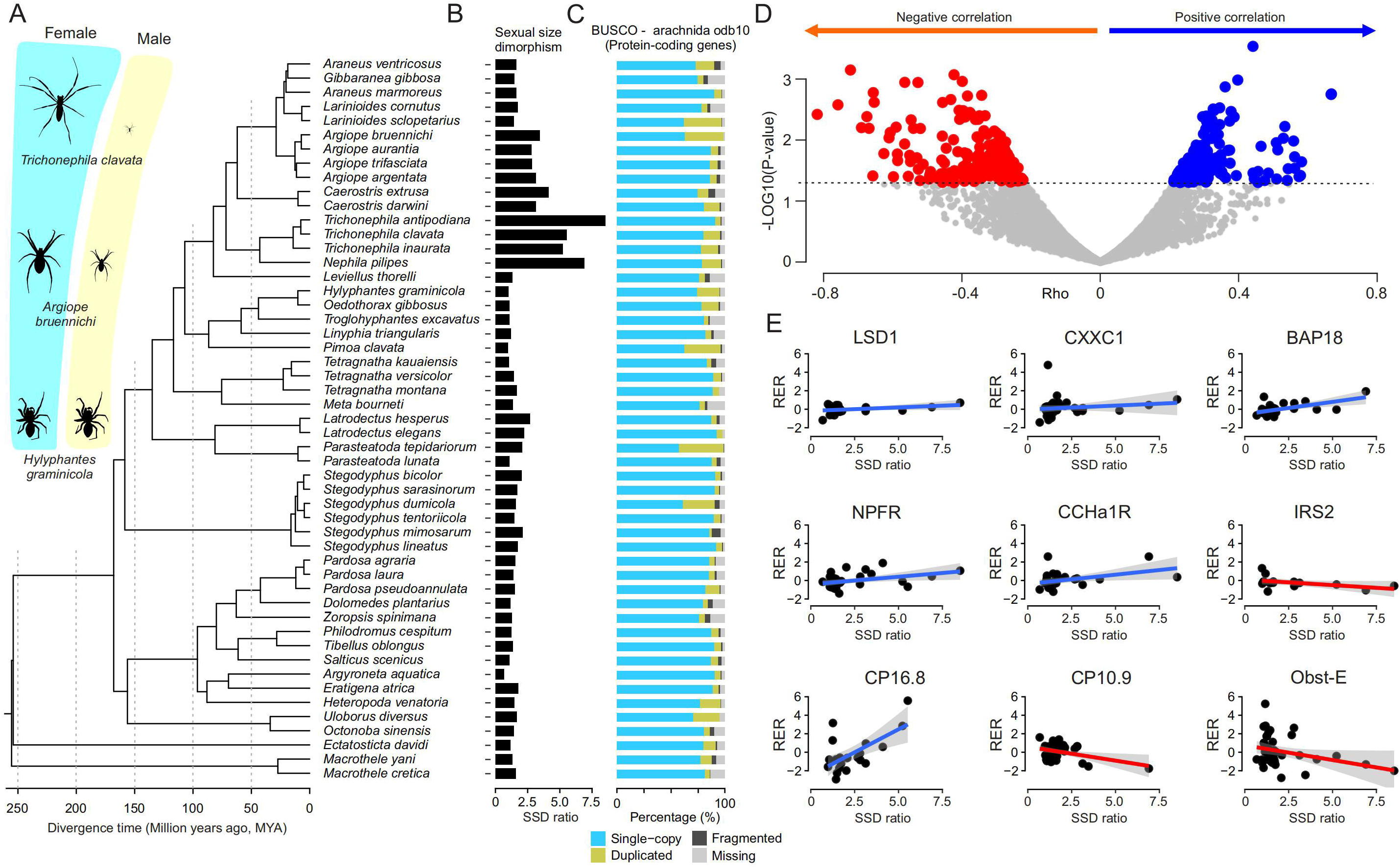
Phylogenomic context and SSD-associated evolutionary-rate shifts across spiders. (A) Phylogeny of the 51 spider species included in the comparative genomic analyses, together with representative female–male body-size differences in selected species, including *Trichonephila clavata*, *Argiope bruennichi*, and *Hylyphantes graminicola*. (B) Sexual size dimorphism (SSD) across the 51 spider species. (C) Completeness of the corresponding protein-coding gene sets assessed using BUSCO with the arachnida_odb10 dataset. (D) Genome-wide associations between relative evolutionary rates (RERs) and SSD estimated using RERconverge. The x-axis shows the correlation coefficient (ρ), and the y-axis shows −log10(P). Genes with nominally significant positive and negative correlations (P < 0.05) are highlighted, whereas nonsignificant genes are shown in gray. The horizontal dashed line indicates P = 0.05. (E) Representative relationships between SSD and RER for selected candidate genes. Regression lines indicate positive or negative associations between RER and SSD.

### Evolutionary-rate shifts in chromatin, growth, and structural genes are associated with increasing SSD

We next asked whether variation in SSD across spiders was associated with changes in protein evolutionary rates. Treating SSD as a continuous trait, we used RERconverge to test associations between gene-specific relative evolutionary rates (RERs) and SSD across the spider phylogeny (Figure 2C; Tables S12 and S13). This analysis identified both positive and negative RER–SSD correlations, representing relative evolutionary acceleration and deceleration, respectively, along branches associated with increasing SSD.

Genes showing nominally significant positive RER–SSD correlations included several chromatin and transcriptional regulators, including *LSD1*, *CXXC1*, and *BAP18*, together with cuticle protein 16.8 (*CP16.8*) and signaling-related genes such as *NPFR* and *CCHa1-R* (Figure 2E). Genes showing negative correlations included growth- and structure-related candidates such as insulin receptor substrate 2-A (*IRS2-A*), cuticle protein 10.9 (*CP10.9*), and *obst-E*. Together, these results indicate that increasing SSD is associated with evolutionary-rate shifts in genes spanning chromatin regulation, growth-related signaling, and cuticular and structural organization.

### Evolutionary rate is associated with expression breadth among male-biased genes

Because SSD represents a sex-specific phenotype, we next examined whether evolutionary-rate variation was related to sex-biased gene-expression patterns in *T. clavata*. We quantified tissue specificity using Tau separately in females and males and tested its relationship with RER within the corresponding sex-biased gene sets (Figure 3).

**Figure 3.**
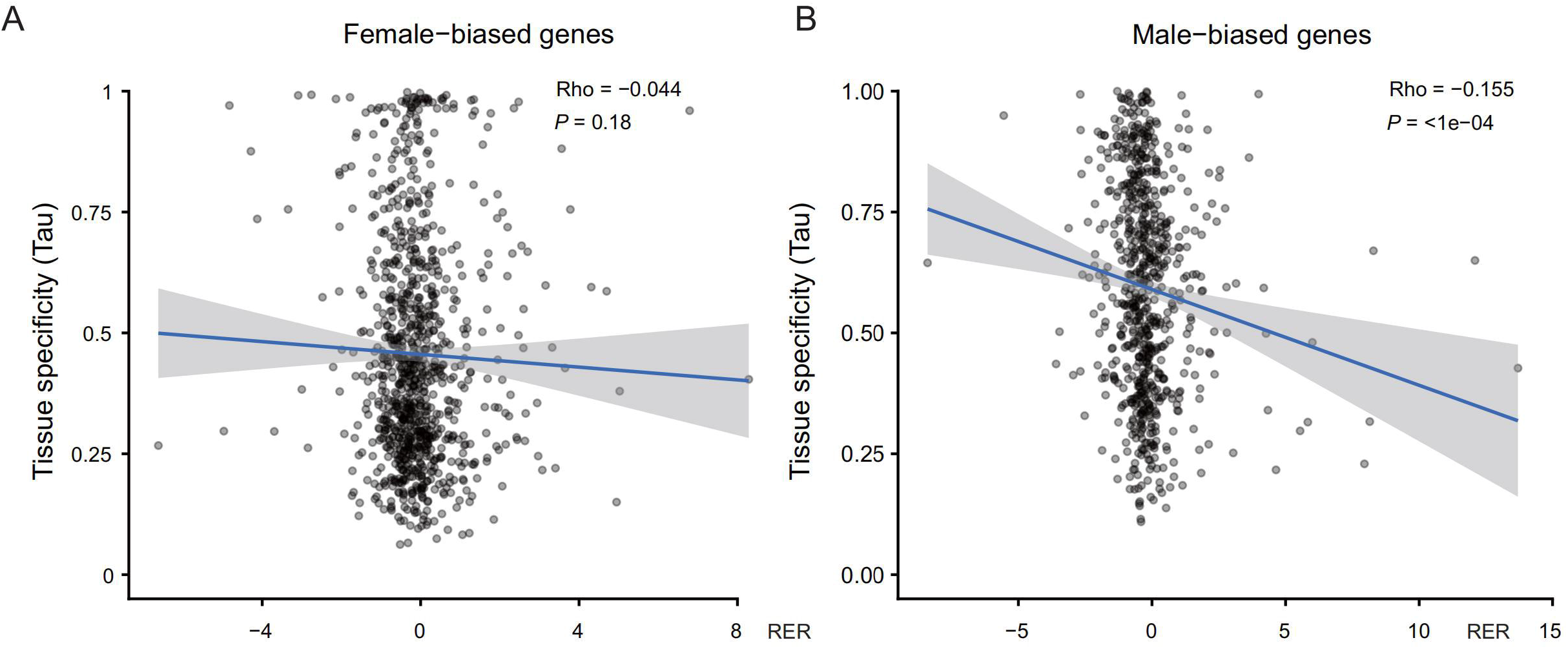
Relationship between tissue specificity and relative evolutionary rate in sex-biased genes. (A) Relationship between relative evolutionary rate (RER) and tissue specificity (Tau) among female-biased genes. (B) Relationship between RER and tissue specificity among male-biased genes.

Among female-biased genes, RER was not significantly associated with female tissue specificity (Spearman’s ρ = −0.044, P = 0.18, n = 941). In contrast, male-biased genes showed a weak but significant negative association between RER and male tissue specificity (ρ = −0.155, P < 10 , n = 711), indicating that male-biased genes with higher relative evolutionary rates tended to be expressed more broadly across male tissues.

We further examined sex-biased genes overlapping with genes showing nominally significant RER–SSD correlations (P < 0.05). Genes with negative RER–SSD correlations were associated with several growth-regulatory and structural systems, including mTOR, PI3K–Akt, AMPK, and Hippo signaling, extracellular matrix–receptor interactions, and cytoskeletal organization. Genes with positive correlations were associated with translation, ribosome biogenesis, RNA processing, biosynthetic metabolism, and neuroactive signaling. Although these pathway enrichments did not remain significant after multiple-testing correction, the gene-level patterns further connected sex-biased expression with growth-, regulatory-, and structural systems highlighted by the comparative evolutionary-rate analysis.

### JHAMT repertoire expansion is positively associated with extreme female-biased SSD

We next asked whether increasing SSD was also associated with changes in gene-family size. Comparative analysis across 51 spider genomes revealed substantial variation in the copy number of juvenile hormone acid O-methyltransferase (JHAMT) genes, with expanded repertoires occurring in several lineages exhibiting extreme female-biased SSD (Figure 4A). Phylogenetic generalized least-squares analysis identified a significant positive association between JHAMT copy number and SSD (Figure 4B), with highly dimorphic orb-weaving lineages, particularly Trichonephila and Nephila, generally possessing larger JHAMT repertoires.

**Figure 4.**
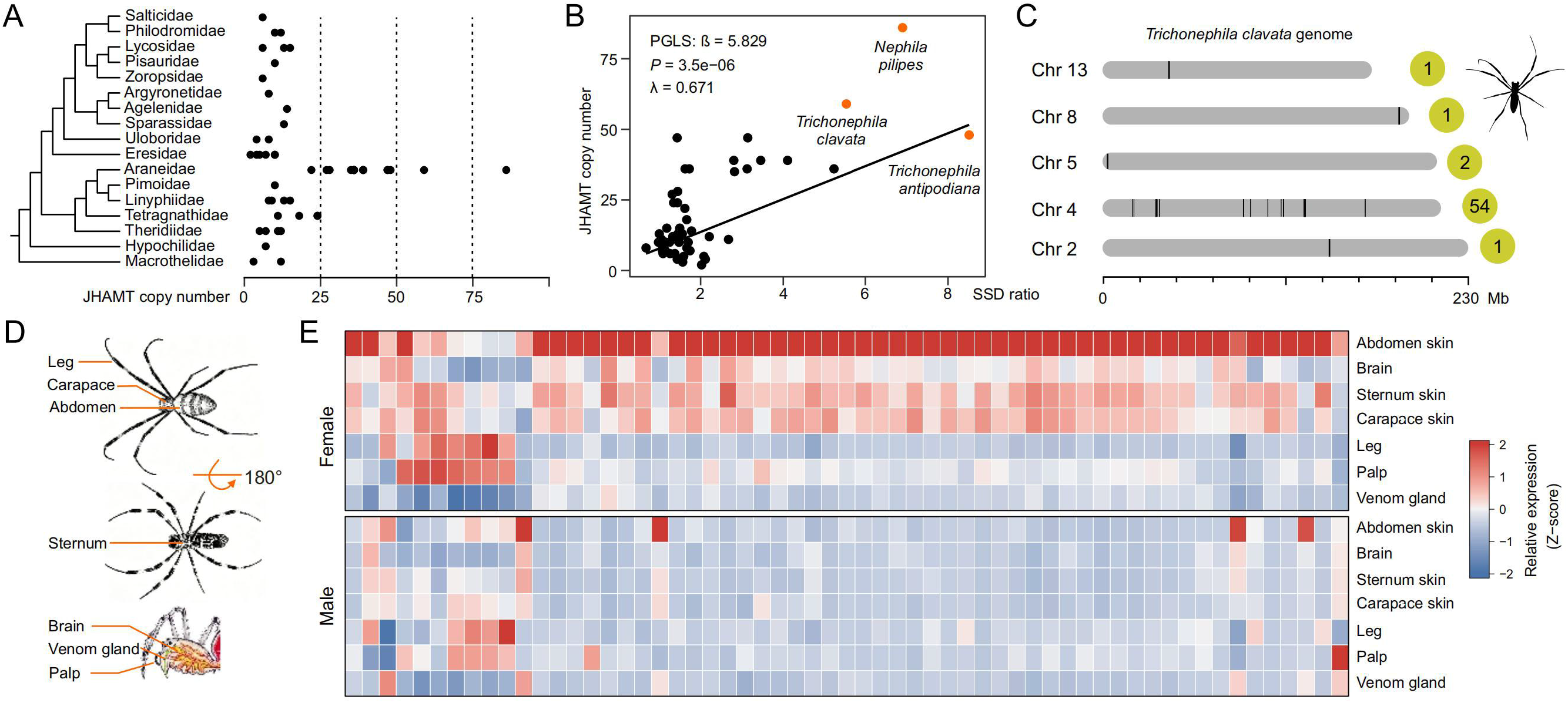
JHAMT repertoire expansion, genomic organization, and expression divergence in spiders. (A) Phylogenetic distribution of JHAMT copy number across sampled spider species. Each point represents the JHAMT copy number of an individual species. (B) Association between JHAMT copy number and sexual size dimorphism (SSD) across spiders evaluated using phylogenetic generalized least squares (PGLS). (C) Chromosomal distribution of 59 annotated JHAMT genes in the *Trichonephila clavata* genome. (D) Schematic showing the seven tissues sampled for transcriptomic profiling: abdominal skin, carapace skin, sternum skin, leg, brain, venom gland, and palp. (E) Heatmap showing JHAMT expression across seven tissues in adult female and male *T. clavata*. Columns represent individual JHAMT genes and rows represent tissues.

To investigate the genomic basis of this expansion, we examined the chromosomal distribution of JHAMT genes in *T. clavata*. JHAMT copies were distributed across four chromosomes, with a prominent cluster on chromosome 4 containing multiple tandemly arranged duplicates (Figure 4C). This genomic organization is consistent with tandem duplication contributing to the expansion of the JHAMT repertoire.

We then asked whether expanded JHAMT paralogs had also diverged in expression. JHAMT genes exhibited pronounced sex-and tissue-dependent expression patterns, with several paralogs showing higher relative expression across female tissues and more restricted expression in males (Figure 4D, 4E). Individual paralogs also differed substantially in tissue specificity. Thus, JHAMT repertoire expansion across spiders is associated with increasing SSD and is accompanied in *T. clavata* by extensive sex- and tissue-specific expression divergence among duplicated genes.

### Female and male *T. clavata* differ in chromatin accessibility and three-dimensional genome organization

The comparative genomic analyses identified SSD-associated changes in genes related to chromatin regulation, growth, and structural organization, together with expansion of the JHAMT repertoire. We therefore asked whether these evolutionary signals were accompanied by sex-specific differences in chromatin organization within *T. clavata*. We profiled genome-wide chromatin accessibility and three-dimensional genome architecture in adult females and males using ATAC-seq and Hi-C.

ATAC-seq revealed extensive sex-specific differences in chromatin accessibility (Figure 5A–C). Accessible chromatin in both sexes was strongly enriched around transcription start sites. More accessible regions were detected in males than in females (229,442 versus 187,407), and male-enriched differentially accessible regions outnumbered female-enriched regions (639 versus 254).

**Figure 5.**
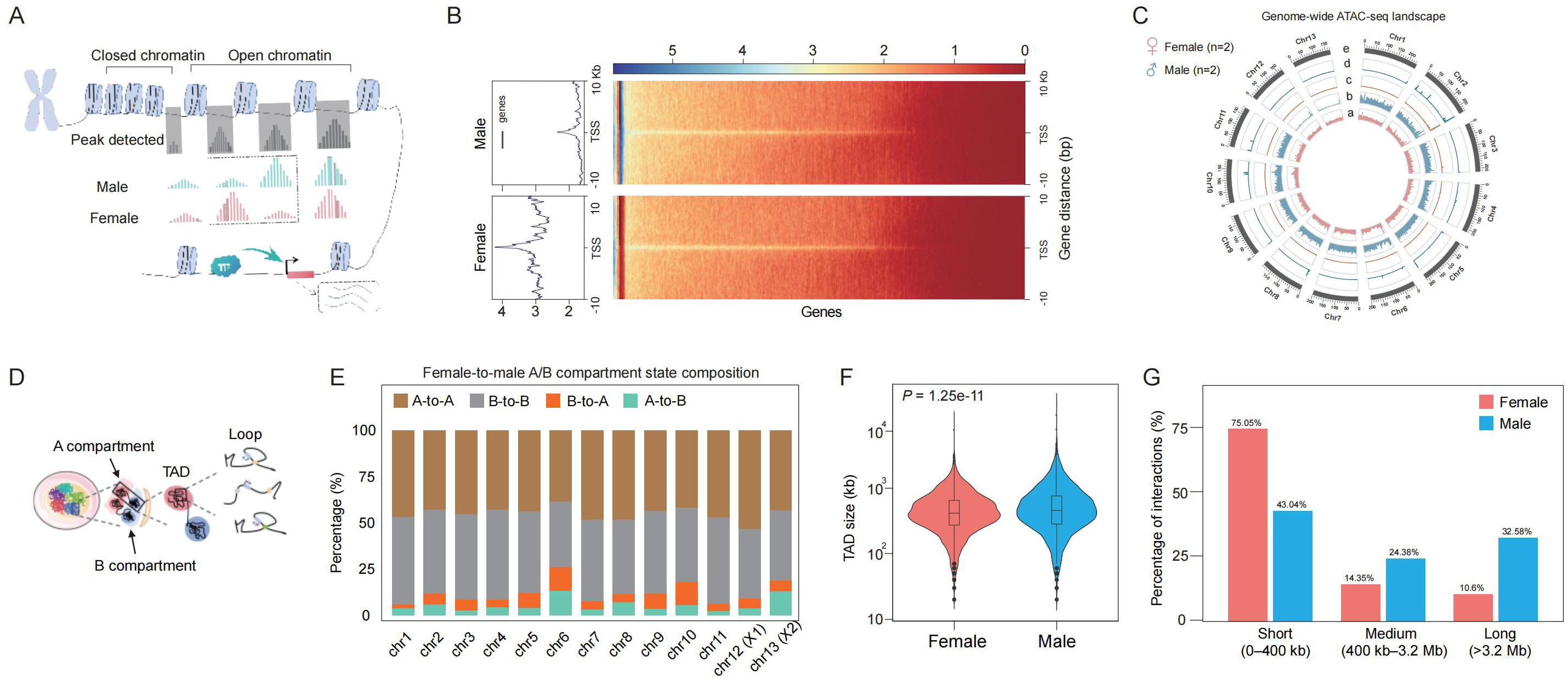
Sex-specific differences in chromatin accessibility and three-dimensional genome organization in adult *Trichonephila clavata*. (A) Schematic representation of ATAC-seq profiling of chromatin accessibility and its relationship to transcriptional regulation. (B) Aggregate ATAC-seq profiles and heatmaps centered on transcription start sites (TSSs) in male (top) and female (bottom) *T. clavata*. Signals are shown across ±10 kb surrounding the TSS. Genes are displayed in the same order and using the same color scale between sexes. (C) Genome-wide landscape of chromatin accessibility in adult female and male *T. clavata*. Circos tracks show (a) female ATAC-seq signals, (b) male ATAC-seq signals, (c) female-enriched differentially accessible regions, (d) male-enriched differentially accessible regions, and (e) chromosome positions. (D) Schematic representation of hierarchical three-dimensional genome organization, including A/B compartments, topologically associating domains (TADs), and chromatin loops. (E) Chromosome-wide comparison of A/B compartment states between adult females and males. Bars indicate the proportions of genomic regions retaining the same compartment state (A-to-A and B-to-B) or switching compartment state (A-to-B and B-to-A) between females and males across individual chromosomes. (F) Distribution of TAD sizes in females and males. Differences were assessed using a Wilcoxon rank-sum test (P = 1.25 × 10 ¹¹). (G) Distribution of cis-interactions according to genomic distance in females and males. Interactions were classified as short-range (0–400 kb), medium-range (400 kb–3.2 Mb), or long-range (>3.2 Mb), with the proportion of each category shown for each sex.

Hi-C analyses further revealed sex-specific differences in higher-order genome organization (Figure 5D– G). More chromatin loops were detected in females than in males (11,772 versus 8,731). A/B compartment comparisons identified both conserved and sex-switched regions, topologically associating domain size distributions differed between the sexes, and cis-interaction frequencies across genomic distances also showed sex-dependent patterns.

Together, these results demonstrate widespread differences between female and male *T. clavata* in both chromatin accessibility and three-dimensional genome organization. These genome-wide differences provided the regulatory context for testing whether endocrine, growth-related, and structural genes also showed coordinated sex-specific molecular divergence.

### Multi-omic integration reveals coordinated sex-specific differences in endocrine, growth-related, and structural genes

We next integrated RNA-seq, ATAC-seq, and Hi-C data to identify genes showing coordinated sex-specific differences across transcriptional and chromatin features. A total of 32 candidate genes showed differential expression, differential chromatin accessibility, and sex-specific chromatin interactions (Figure S13; Table S15). These genes were associated with endocrine, metabolic, signaling, cytoskeletal, and structural functions.

Genes supported by both differential expression and sex-specific chromatin interactions were enriched for hormone- and steroid-related metabolic processes, including isoprenoid and terpenoid metabolism (Figure 6). Sex-biased transcription was also observed among genes associated with endocrine signaling, epidermal development, and structural organization.

**Figure 6.**
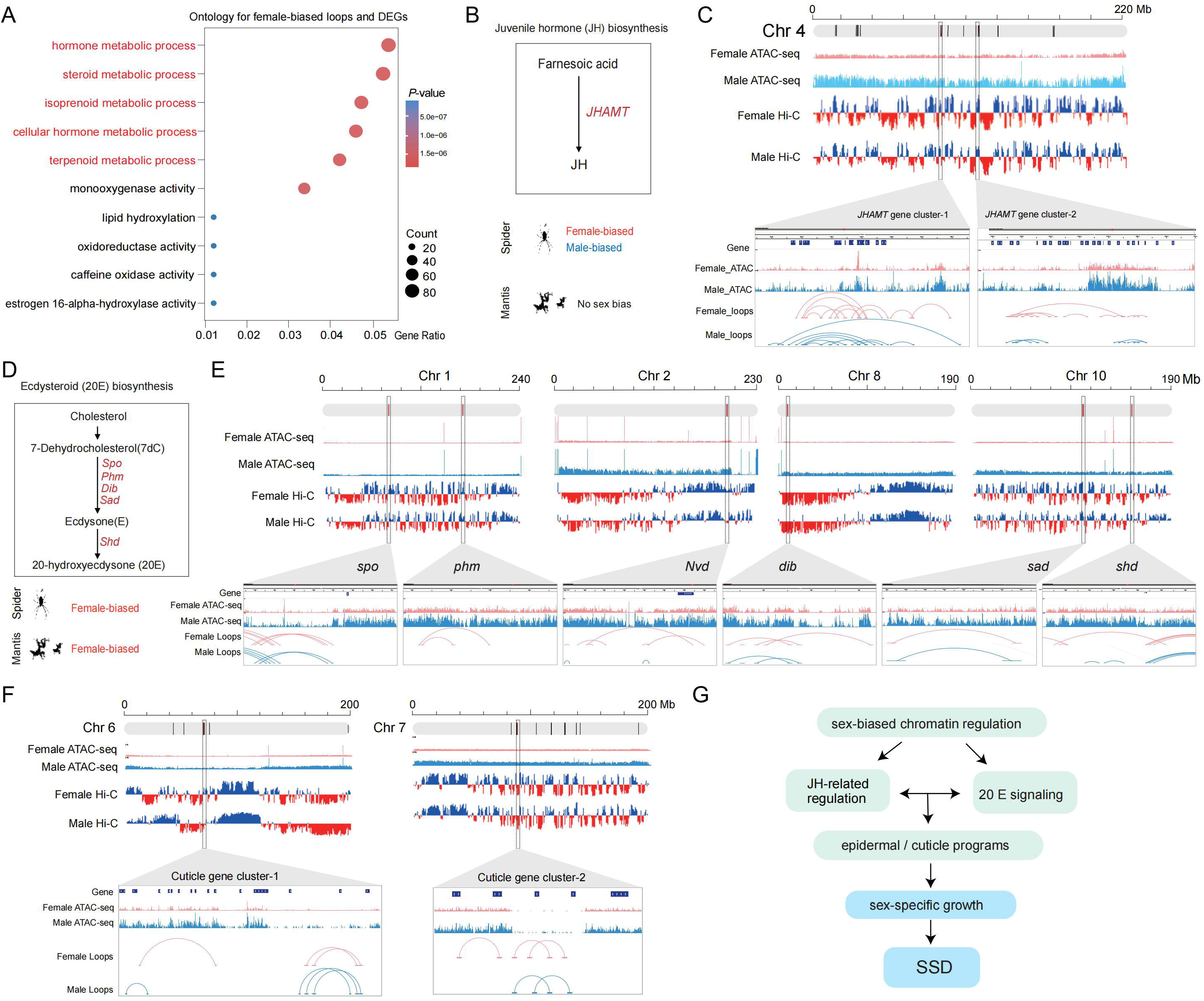
Multi-omic integration reveals coordinated sex-specific differences in endocrine, growth-related, and structural genes in *Trichonephila clavata*. (A) Gene Ontology (GO) enrichment analysis of genes associated with female-specific chromatin loops and differential expression. (B) Simplified juvenile hormone-related biosynthetic pathway highlighting JHAMT. (C) Chromatin accessibility and local chromatin-interaction profiles at representative JHAMT gene clusters. (D) Schematic representation of the ecdysteroid (20E) biosynthetic pathway. (E) Chromatin accessibility and local chromatin-interaction profiles at representative ecdysteroid-biosynthetic genes, including *spo*, *phm*, *nvd*, *dib*, *sad*, and *shd*. (F) Chromatin accessibility and local chromatin-interaction profiles at representative cuticular and structural gene clusters. (G) Schematic summary of candidate endocrine, growth-related, epidermal, and cuticular systems associated with sex-specific molecular divergence and extreme SSD.

The expanded JHAMT repertoire provided a direct connection between the comparative genomic and within-species multi-omic analyses. In addition to their sex- and tissue-biased expression, JHAMT loci on chromosome 4 exhibited pronounced sex-specific differences in chromatin accessibility and local chromatin interactions (Figure 6). Thus, the positive association between JHAMT copy number and SSD across spiders was accompanied in *T. clavata* by sex-specific transcriptional and chromatin variation at the expanded loci.

Similar patterns extended beyond JHAMT. Putative orthologs of ecdysteroid-biosynthetic genes, including *nvd*, *spo*, *Phm*, *dib*, *sad*, and *shd*, showed sex-specific differences in transcription, chromatin accessibility, and/or local chromatin interactions (Figure 6). Coordinated differences were also detected in genes associated with insulin and growth regulation and in epidermal, cuticular, and other structural processes.

Together, the multi-omic analyses reveal coordinated sex-specific differences in transcription, chromatin accessibility, and chromatin interactions across endocrine, growth-related, epidermal, and cuticular genes. These within-species regulatory patterns converge on several of the same biological systems highlighted by the comparative genomic analyses, linking SSD-associated evolutionary changes across spiders with sex-specific molecular divergence in an extremely dimorphic species.

## Discussion

Our study identifies evolutionary and regulatory changes associated with extreme female-biased sexual size dimorphism (SSD) in spiders across several levels of genomic variation. Comparative analyses across 51 spider species revealed SSD-associated shifts in protein evolutionary rates and gene-family size, whereas sex-resolved multi-omic profiling in *Trichonephila clavata* further identified coordinated differences in transcription and chromatin regulation between females and males. These patterns converge on biological systems related to growth regulation, endocrine signaling, chromatin regulation, and cuticular and structural development. Together, our results suggest that extreme SSD is associated not with a single genomic change, but with evolutionary and regulatory divergence distributed across multiple molecular processes.

### SSD-associated evolutionary-rate variation highlights growth, chromatin, and structural systems

The comparative genomic analyses identified evolutionary-rate shifts associated with increasing SSD in genes involved in growth regulation, chromatin-related functions, and structural organization. Among genes showing negative RER–SSD correlations, *IRS2-A* links SSD-associated rate variation to growth-related signaling, whereas *CP10.9* and *obst-E* implicate cuticular and structural processes. These patterns are consistent with the established importance of nutrient- and growth-sensing pathways in coordinating developmental growth in arthropods (Callier and Nijhout, 2013; Koyama and Mirth, 2018). Positive correlations were also detected in chromatin and transcriptional regulators, including *LSD1*, *CXXC1*, and *BAP18*, together with additional signaling and structural genes. Thus, the evolutionary-rate signals associated with SSD extend across molecular systems that could plausibly influence growth, tissue development, and the structural consequences of prolonged or differential growth.

The involvement of chromatin-related genes is particularly notable because changes in chromatin regulation can influence the transcriptional deployment of developmental programs without requiring wholesale changes in pathway composition. However, RER correlations alone do not distinguish whether the observed rate shifts reflect altered selective constraint, adaptive evolution, or other changes in sequence evolution. Likewise, because pathway-level enrichments did not remain significant after multiple-testing correction, these results are best interpreted as gene-level associations rather than evidence for coordinated pathway-wide evolutionary shifts.

Sex-biased expression provides an additional context for these evolutionary patterns. Among male-biased genes, higher RER was weakly associated with broader expression across male tissues, whereas no significant relationship was detected among female-biased genes. Although the biological significance of this difference remains uncertain, it suggests that evolutionary-rate variation in the male-biased gene set is not restricted to narrowly tissue-specific genes. More broadly, the overlap between sex-biased expression and SSD-associated RER signals places sequence evolution and sex-specific transcription within a shared set of growth-related and regulatory functions.

### JHAMT repertoire expansion links gene-family evolution with extreme SSD

The positive association between JHAMT copy number and SSD identifies gene-repertoire evolution as a second major dimension of genomic variation associated with extreme dimorphism. JHAMT is a key component of juvenile-hormone biosynthesis in insects (Shinoda and Itoyama, 2003), whereas studies in the wolf spider *Pardosa pseudoannulata* have implicated multiple JHAMT homologs and methyl farnesoate in juvenile growth and molting (Yang et al., 2021a, 2022). These observations support a developmental role for JHAMT-related endocrine biology in spiders while also indicating that spider endocrine regulation differs from the canonical insect juvenile-hormone system.

The genomic organization of JHAMT in *T. clavata* further suggests that tandem duplication contributed to repertoire expansion. Gene duplication can provide opportunities for ancestral functions to be partitioned among paralogs or for new regulatory and biochemical properties to evolve (Force et al., 1999; Innan and Kondrashov, 2010). Consistent with this possibility, expanded JHAMT paralogs in *T. clavata* exhibited pronounced sex- and tissue-biased expression, indicating substantial regulatory divergence among duplicated copies. These expression differences do not establish biochemical neofunctionalization, but they suggest that JHAMT expansion was accompanied by diversification in transcriptional deployment.

JHAMT therefore provides the clearest connection between the comparative and within-species components of our study. Across spiders, larger JHAMT repertoires were associated with greater SSD, whereas in *T. clavata*, duplicated JHAMT loci also showed sex-specific transcriptional and chromatin differences. This combination of gene-family expansion, expression divergence, and chromatin variation identifies JHAMT as a particularly informative candidate system for testing how endocrine gene-repertoire evolution relates to sex-specific growth. A similar trend observed in sampled mantises is intriguing, but broader comparative sampling will be required before inferring convergence or a shared mechanism across arthropod lineages.

### Sex-specific transcriptional and chromatin divergence converges on endocrine, growth-related, and structural genes

The evolutionary associations involving chromatin-related genes motivated us to examine whether extreme SSD was also accompanied by regulatory divergence between females and males in *T. clavata*. Genome-wide analyses revealed differences in chromatin accessibility, chromatin loops, compartment states, and domain organization. More accessible regions were detected in males, whereas more chromatin loops were detected in females. These contrasting patterns indicate sex-specific differences in distinct aspects of chromatin organization, but neither feature count alone implies globally greater regulatory activity in either sex.

Integrating RNA-seq, ATAC-seq, and Hi-C extended these genome-wide observations to specific biological systems. A set of candidate genes showed coordinated differences in transcription, chromatin accessibility, and chromatin interactions between the sexes. Importantly, these genes were not restricted to a single regulatory pathway. Instead, the multi-omic signals converged on endocrine, growth-related, epidermal, cuticular, and other structural functions.

JHAMT again provided the clearest example of this convergence. Expanded JHAMT loci showed sex- and tissue-biased transcription together with sex-specific chromatin accessibility and local chromatin interactions. Putative ecdysteroid-related genes also exhibited sex-dependent transcriptional and chromatin patterns. Independent functional studies in *P. pseudoannulata* have shown that ecdysteroid signaling influences spider development and molting (Yang et al., 2021b, 2022), supporting the biological plausibility of these genes as candidates while not establishing equivalent functions in *T. clavata*.

Comparable sex-specific molecular differences were also observed in genes involved in insulin and growth regulation and in epidermal and cuticular processes. These functional categories are biologically connected through arthropod growth and molting, in which endocrine signals coordinate tissue growth, epidermal remodeling, and cuticle production (Charles, 2010; Yamanaka et al., 2013). Their repeated appearance across comparative genomic and multi-omic analyses therefore raises the possibility that extreme SSD involves coordinated divergence among endocrine, growth-regulatory, and structural systems rather than changes confined to a single pathway.

At the same time, the present data do not establish a causal chain linking sequence evolution, gene duplication, chromatin differences, transcription, and body-size divergence. Chromatin accessibility and three-dimensional contacts provide regulatory context, but differences in these features do not by themselves demonstrate direct effects on gene expression. Likewise, the co-occurrence of evolutionary and regulatory signals in the same functional systems does not establish that these processes jointly generate SSD.

An additional limitation is that most molecular data were obtained from adults. Our analyses therefore characterize molecular features associated with established SSD rather than the developmental processes through which female and male growth trajectories diverge. Tissue-specific RNA-seq and whole-individual ATAC-seq and Hi-C also differ in cellular composition, limiting direct cross-omic interpretation. High sequence similarity among duplicated genes may further affect paralog-specific expression and chromatin estimates. Future analyses of matched tissues across juvenile development, together with direct measurements of endocrine activity and functional perturbation of candidate genes or regulatory elements, will be necessary to determine which of these associations contribute to sex-specific growth.

## Conclusion

In summary, our results connect extreme female-biased SSD in spiders with evolutionary-rate shifts, endocrine gene-repertoire expansion, and sex-specific transcriptional and chromatin divergence. Rather than identifying a single molecular driver, our findings reveal recurring associations involving endocrine, growth-regulatory, chromatin-related, epidermal, and cuticular systems. The convergence of comparative genomic and multi-omic signals on these functions provides a framework for understanding the evolutionary and regulatory changes associated with extreme body-size dimorphism in spiders.

## Method and Materials

### Study design

We compiled adult female and male body-size data for a large number of spider species spanning a broad range of sexual size dimorphism (SSD) and analyzed their genomes to investigate the evolutionary genomic basis of extreme SSD. We used comparative genomic analyses to examine associations between SSD evolution, gene family evolution, and evolutionary-rate shifts. We further focused on the extremely female-biased Joro spider, *Trichonephila clavata*, and generated parallel RNA-seq data from seven tissues of adult females and males, together with ATAC-seq and Hi-C datasets from both sexes. These datasets were integrated to investigate sex-specific differences in gene expression, chromatin accessibility, and three-dimensional genome organization associated with extreme SSD.

### Phenotypic data collection

To systematically compare the extent of sexual size dimorphism across arthropods, we compiled publicly available adult female and male body-length data for 23 representative arthropod species from the published literature and open-access databases (Supplementary Table S1). Similarly, we assembled a dataset of sex-specific adult body lengths for 51 spider species with publicly available genome assemblies (Supplementary Table S2). We included only body-length measurements from sexually mature individuals. When body length was reported as a range, we used the midpoint of the reported minimum and maximum values. All measurements were standardized to millimetres. Sexual size dimorphism (SSD) was calculated as:

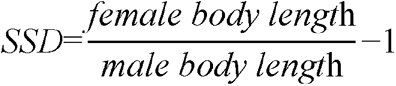

Species with |SSD| ≤ 0.10 were classified as exhibiting no pronounced sexual size dimorphism, whereas species with SSD > 0.10 were classified as female-biased, whereas those with SSD < −0.10 were classified as male-biased, respectively. Continuous SSD values were retained for subsequent correlation and phylogenetic comparative analyses.

### Genomic data collection

We collected publicly available spider genome assemblies from online database, including NCBI Genome (https://www.ncbi.nlm.nih.gov/genome), ScienceDB (https://www.scidb.cn/en), and GigaDB (http://gigadb.org/), as of December 2025. Further, we assessed genome assembly completeness assessment of genome assemblies using BUSCO v5.4.5 (Manni et al., 2021) with arachnida_odb10 data set (N = 2,934). When multiple genome assemblies were available for a spider species, the most complete and/or updated assembly (i.e. chromosome-level) was selected (Supplementary Table S2). We downloaded the genome of Macrothele yani (GCA_039090855.1), and Macrothele cretica (GCA_964417605.1) as outgroup for phylogenomic analysis . We extracted completed single-copy BUSCO genes from each spider genome, and further assembled a phylogenomic dataset including 51 spider species. In addition, we performed protein sequence alignments using MAFFT v7.515 (Katoh and Standley, 2013), and trimmed the alignments using trimAl v1.4 (Capella-Gutiérrez et al., 2009). We assembled a concatenated dataset that included all 1:1 BUSCO single-copy genes with a minimum length of 200 amino acids. Finally, we used ModelFinder (Kalyaanamoorthy et al., 2017) to determine the best-fit model of sequence evolution (protein model: LG + I + G4) and constructed the maximum likelihood (ML) phylogenetic tree using IQ-TREE2 (Minh et al., 2020) with 1,000 bootstrap replicates.

### Molecular dating

Divergence times were estimated for the 51 spider species using MCMCTree in PAML (Yang, 2007). The species tree topology was fixed, with fossil calibrations applied to the Nephilinae stem (43.0–47.8 Ma) and Araneoidea stem (125–135 Ma) (Magalhaes et al., 2020). The root age was constrained to a maximum of 374 Ma. Independent MCMC runs were performed to assess convergence.

### Ortholog identification

In addition to the BUSCO gene set described above, which included only curated single-copy orthologs from OrthoDB database (https://www.orthodb.org), we used FastOMA (Majidian et al., 2025) to extend and explore gene orthology across 51 spider genomes following the pipeline described in our recent study (Tong et al., 2026).

### Gene family evolution

Building on the orthologous groups identified by FastOMA, we used CAFE5 (Mendes et al., 2020) to characterize gene family expansion and contraction across spider genomes. We first generated a gene-count matrix based on the number of genes assigned to each orthologous family in each species. Together with an ultrametric species tree, this matrix was used as input for CAFE5, which models gene family gain and loss under a stochastic birth–death process and estimates lineage-specific changes in family size. Gene families showing significant expansions or contractions were identified using a significance threshold of P < 0.05. We examined whether variation in gene family size was associated with the evolution of SSD across spiders.

We sought to identify gene families whose expansion was significantly positively associated with increasing SSD across spider species. Among these families, we focused on the juvenile hormone acid O-methyltransferase (JHAMT) gene family because of its key role in juvenile hormone biosynthesis and growth regulation. Specifically, we identified candidate JHAMT homologs across spider genomes using MMseqs2 v13.45111 (Steinegger and Söding, 2017) with known JHAMT protein sequences as queries. Four iterative rounds of blastp-like similarity searches were performed using an E-value threshold of 1 × 10^-3^ to recover both closely and more distantly related homologs. We then examined the candidate proteins with InterProScan (Jones et al., 2014) based on Pfam annotations (Mistry et al., 2021) and retained only sequences containing the characteristic methyltransferase domain. After removing redundant protein isoforms and candidates lacking the expected domain architecture, we defined JHAMT copy number for each spider species as the number of validated genes located at distinct genomic loci. For *T. clavata*, we retrieved the genomic coordinates of validated JHAMT copies from the genome annotation and visualized their chromosomal distribution using MG2C (Chao et al., 2021).

### Assessment of the association between evolutionary rate changes and SSD evolution

To test whether the shifts in evolutionary rates are associated with the evolution of SSD, we computed the relative evolutionary rate (RER) for each spider species at gene level. Specifically, we performed protein sequence alignments of shared orthologs across all 51 species using MAFFT (Nakamura et al., 2018). Next, we inferred a gene tree and estimated branch length for each shared ortholog using R package Phangorn (Schliep, 2011). We calculated RER for each phylogenetic node on gene tree using R package, RERconverge (Kowalczyk et al., 2019). Further, we tested for the associations between branch-specific RER and SSD changes using *correlateWithContinuousPhenotype*, which calculates Pearson correlations across phylogenetic branches. To reduce the influence of extreme values, both RERs and trait values were winsorized prior to correlation analysis. Genes with positive correlation coefficients were interpreted as showing accelerated relative evolutionary rates along branches associated with increasing SSD, whereas negative coefficients indicated relative evolutionary deceleration.

### Study species and sample collection

We collected sexually mature female and male *T. clavata* individuals from the same sampling locality in Dali, Yunnan, China in September, 2025. To minimize potential confounding effects associated with developmental stage, sampling location, and collection time, we included only mature individuals collected under these standardized conditions.

For transcriptomic profiling by RNA sequencing (RNA-seq), we dissected seven tissues from female and male spiders, including brain, abdominal skin, carapace skin, sternum skin, leg, palp, and venom gland. We generated three biological replicates for each sex–tissue combination, with each replicate consisting of pooled tissue samples from three adult individuals of the same sex. Immediately after dissection, we transferred all tissue samples into RNase-free 1.5 mL microcentrifuge tubes, flash-froze them in liquid nitrogen, and stored at −80 °C until RNA extraction and sequencing.

For assay for transposase-accessible chromatin using sequencing (ATAC-seq), we generated chromatin accessibility profiles from whole body samples of four adult individuals, including two females and two males. We immediately flash-froze the samples in liquid nitrogen and stored them at −80 °C.

For high-throughput chromosome conformation capture (Hi-C) analysis, we generated new Hi-C data from whole body samples of three adult individuals, including two males (Tclav-M-Hic-1 and Tclav-M-Hic-2) and one female (Tclav-F-Hic-1). We immediately flash-froze the samples in liquid nitrogen and stored them at −80 °C until further processing. An additional female Hi-C dataset (Tclav-F-Hic-2) was obtained from the Genome Sequence Archive (GSA) of the National Genomics Data Center (NGDC), under BioProject accession PRJCA014503 (Hu et al., 2023). Thus, two female and two male Hi-C datasets were included in the subsequent analyses.

### RNA-sequencing and comparative analysis

We extracted total RNA using TRIzol reagent (Invitrogen, USA) according to the manufacturer’s instructions. We prepared RNA-seq libraries using the NEBNext Ultra RNA Library Prep Kit for Illumina (New England Biolabs, USA) and sequenced them on an Illumina NovaSeq 6000 platform. We filtered raw reads using BBTools v38.67 (Bushnell, 2014) and aligned the clean reads to the *T. clavata* reference genome using HISAT2 v2.2.1 (Kim et al., 2019). We quantified gene-level read counts using featureCounts v2.0.3 (Liao et al., 2014), retaining uniquely mapped and properly paired reads. We calculated transcripts per million (TPM) values for expression-level comparisons and visualization, whereas we used raw read counts for differential expression analysis.

We performed differential expression analysis between females and males separately for each tissue using R package DESeq2 (Love et al., 2014). We defined the contrast as female versus male, such that positive log2 fold-change values indicated higher expression in females and negative values indicated higher expression in males. We considered genes with |log2 fold change| > 1 and a Benjamini–Hochberg-adjusted P < 0.05 to be significantly sex-biased. Genes with significant positive fold changes were classified as female-biased, whereas those with significant negative fold changes were classified as male-biased. For downstream analyses, we combined female-biased genes identified across the seven tissues into a global female-biased gene set and similarly combined male-biased genes into a global male-biased gene set. Genes showing opposite directions of sex bias among different tissues were retained in both corresponding gene sets.

We quantified tissue specificity of gene expression using the Tau (τ) metric implemented in tspex (Camargo et al., 2020) and analyzed females and males separately. For each sex, we averaged DESeq2-normalized expression values across three biological replicates within each tissue and excluded genes with no detectable expression across all seven tissues. Tau values range from 0 to 1, with values approaching 1 indicating highly tissue-specific expression and values approaching 0 indicating broadly distributed expression across tissues.

To examine the relationship between evolutionary rate and sex-specific expression patterns, we matched sex-biased genes to their gene-specific RER estimated by RERconverge. For female-biased genes, we tested the association between RER and female-specific Tau values, whereas for male-biased genes, we tested the association between RER and male-specific Tau values. We retained only genes with both RER and Tau estimates and evaluated the associations using Spearman’s rank correlation.

### ATAC-seq library preparation and data analysis

We generated ATAC-seq libraries from four sexually mature *T. clavata* individuals, including two females and two males. Library preparation followed the standard ATAC-seq protocol with minor modifications (Buenrostro et al., 2015). Briefly, nuclei were isolated from each sample and subjected to Tn5 transposase-mediated tagmentation, followed by DNA purification, library amplification, and quantification. Finally, all libraries were sequenced in the same lane on an Illumina NovaSeq 6000 platform (Frasergen Co., Ltd., Wuhan, China).

We assessed the quality of raw sequencing reads using FastQC v0.12.1 (Andrews, 2010) and removed adapter sequences and low-quality bases using Trimmomatic v0.39 (Bolger et al., 2014). We aligned the filtered reads to the *T. clavata* reference genome using Bowtie2 v2.5.2 (Langmead and Salzberg, 2012). We removed low-quality alignments, PCR duplicates, and reads mapped to organellar genomes before downstream analyses. We evaluated ATAC-seq library quality based on fragment-size distributions and enrichment of sequencing signals around transcription start sites (TSSs), with TSS enrichment profiles generated using deepTools v3.5.4 (Ramírez et al., 2016).

We identified accessible chromatin regions using MACS2 v2.2.7.1 (Zhang et al., 2008) and annotated the resulting peaks relative to genomic features using ChIPseeker v1.38.0 (Yu et al., 2015). To identify sex-biased chromatin accessibility, we compared ATAC-seq signals between females and males using DiffBind (Stark and Brown, 2011), with differential testing performed using R package, DESeq2. Regions with |log2 fold change| > 0.58 and a false discovery rate (FDR) < 0.05 were defined as differentially accessible regions (DARs). DARs with positive fold changes were considered female-enriched, whereas those with negative fold changes were considered male-enriched.

To investigate potential transcriptional regulators underlying sex-biased chromatin accessibility, we performed motif enrichment analyses separately for female- and male-enriched DARs using the MEME Suite (Bailey et al., 2015), with transcription-factor motifs obtained from the JASPAR database (Rauluseviciute et al., 2024). We further examined transcription-factor occupancy patterns using RGT-HINT v1.0.2 (Li et al., 2019) and performed differential footprinting analyses between females and males to identify transcription factors showing sex-specific binding signatures.

### Hi-C library preparation and data analysis

We generated in situ Hi-C libraries from two adult male and one adult female *T. clavata* individuals collected in this study. Library construction was performed following an in situ Hi-C protocol adapted from Belaghzal et al. (2017). Briefly, we cross-linked chromatin, digested genomic DNA with MboI, filled the resulting DNA ends with biotin-labeled nucleotides, and performed proximity ligation to capture spatially adjacent chromatin fragments. After reversing the cross-links, we purified the ligated DNA and prepared sequencing libraries. We sequenced the libraries on an Illumina NovaSeq 6000 platform using 150-bp paired-end reads.

For comparative analyses, we aligned clean paired-end reads to the *T. clavata* reference genome using Bowtie2 v2.5.2 (Langmead and Salzberg, 2012). We processed mapped reads using HiC-Pro (Servant et al., 2015) to assign read pairs to MboI restriction fragments and remove invalid ligation products, multimapped reads, singletons, and PCR duplicates. Only valid interaction pairs between distinct restriction fragments were retained for downstream analyses. We then generated Hi-C contact matrices using Juicer v1.6 (Durand et al., 2016) at 20-, 40-, 150-, and 500-kb resolutions and normalized the matrices using Knight–Ruiz (KR) matrix balancing (Knight and Ruiz, 2013). We visualized chromosome-scale interaction maps using HiCExplorer v3.6 (Wolff et al., 2020).

We identified A/B compartments from 500-kb KR-normalized contact matrices using principal component analysis implemented in Juicer Tools v1.6. We used the first principal component (PC1) to assign compartment states and determined the orientation of the eigenvector based on transcription start site density, with regions enriched for transcription start sites assigned to the active A compartment. We identified topologically associating domain (TAD) boundaries using insulation-score analysis (Crane et al., 2015).

We detected chromatin loops using HiCCUPS implemented in Juicer Tools at 5-, 10-, and 20-kb resolutions with KR-normalized contact matrices (Rao et al., 2014). We retained loops with a false discovery rate (FDR) ≤ 0.05 and excluded low-confidence interactions supported by no more than two read pairs. The resulting compartment, TAD, and loop annotations were subsequently used to compare three-dimensional genome organization between females and males.

### Multi-omics integration and visualization

We integrated RNA-seq, ATAC-seq, and Hi-C datasets to identify genes showing coordinated sex-specific differences across transcriptional, chromatin-accessibility, and three-dimensional chromatin organization. We jointly examined differential gene expression, differential chromatin accessibility, and sex-specific chromatin-loop interactions to prioritize genes supported by multiple regulatory features. Genes showing concordant evidence across these datasets were considered multi-omic candidate genes and were further evaluated for their potential associations with sexual size dimorphism. Finally, we visualized representative genomic regions using IGV v2.19.1 (Robinson et al., 2011), including gene annotations, RNA-seq expression profiles, ATAC-seq accessibility signals, and Hi-C chromatin interactions. Gene-expression heatmaps were generated using TBtools-II (Chen et al., 2023).

### Gene ontology and pathway enrichment analysis

We performed Gene Ontology (GO) and Kyoto Encyclopedia of Genes and Genomes (KEGG) enrichment analyses to characterize the biological functions and pathways associated with candidate genes identified from different genomic and multi-omic analyses. Gene sets derived from RERconverge, gene family evolution, differential gene expression, differential chromatin accessibility, Hi-C-based chromatin interactions, and multi-omics integration were analyzed separately using R package clusterProfiler (Wu et al., 2021). Enrichment significance was assessed using the Benjamini-Hochberg procedure, and terms or pathways with an adjusted P < 0.05 were considered significantly enriched. Enriched GO terms and KEGG pathways were subsequently compared across analyses to identify shared and analysis-specific biological processes potentially associated with the evolution and regulation of sexual size dimorphism.

## Data availability

The data sequenced in this manuscript were submitted in scienceDB dataset (https://cstr.cn/31253.11.sciencedb.012wk).

## Acknowledgments

This work was supported in part by the High Performance Computing (HPC) clusters at Southwest University. This research is supported by the National Natural Science Foundation of China (32570539), the Key Project of Chongqing Municipality (cstc2019jcyj-zdxmX0006), the Science & Technology Fundamental Resources Investigation Program (2024FY100403), and the Fund on survey of Invertebrates from Yintiaoling Nature Reserve (CQS24C00333) by Zhi-Sheng Zhang.

## Author contributions

Z.F. and C.T. wrote the manuscript, prepared the figures, and performed data analyses. Z.F. and J.G. performed initial data analyses. L.W., T.R., W.P., W.W., and J.K. participated in sample collection and data curation. Y.H. revised the manuscript. C.T. and Z.Z. designed the study. All authors revised and approved the final manuscript.

