## Supplemental figures 1-13 for "Evolutionary and multi-omic divergences associated with the extreme sexual size dimorphism in spiders": supplementary Figures.pdf

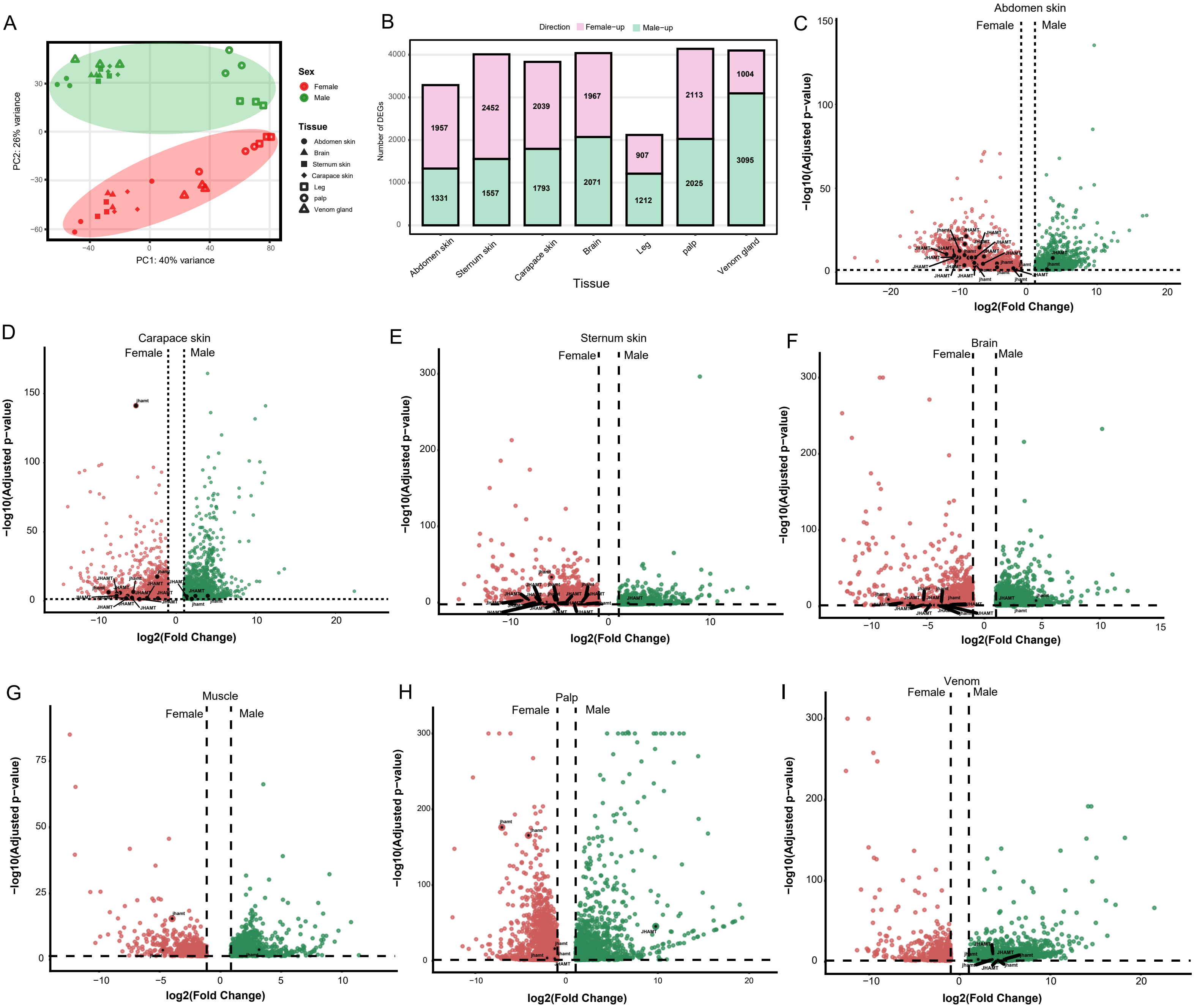


**Figure S1. Sex-biased gene expression across adult tissues of *Trichonephila clavata*.**
(A) Principal component analysis (PCA) of RNA-seq samples from females and males across seven tissues . Colors indicate sex and shapes indicate tissue type. (B) Numbers of female-upregulated and male-upregulated differentially expressed genes (DEGs) in each tissue.(C – I) Volcano plots showing sex-biased gene expression in abdomen skin (C), carapace skin (D), sternum skin (E), brain (F), muscle (G), palp (H), and venom gland (I). Red and green points indicate female- and male-upregulated genes, respectively. Dashed lines indicate the thresholds used to define significant differential expression. Selected JHAMT genes are labeled.

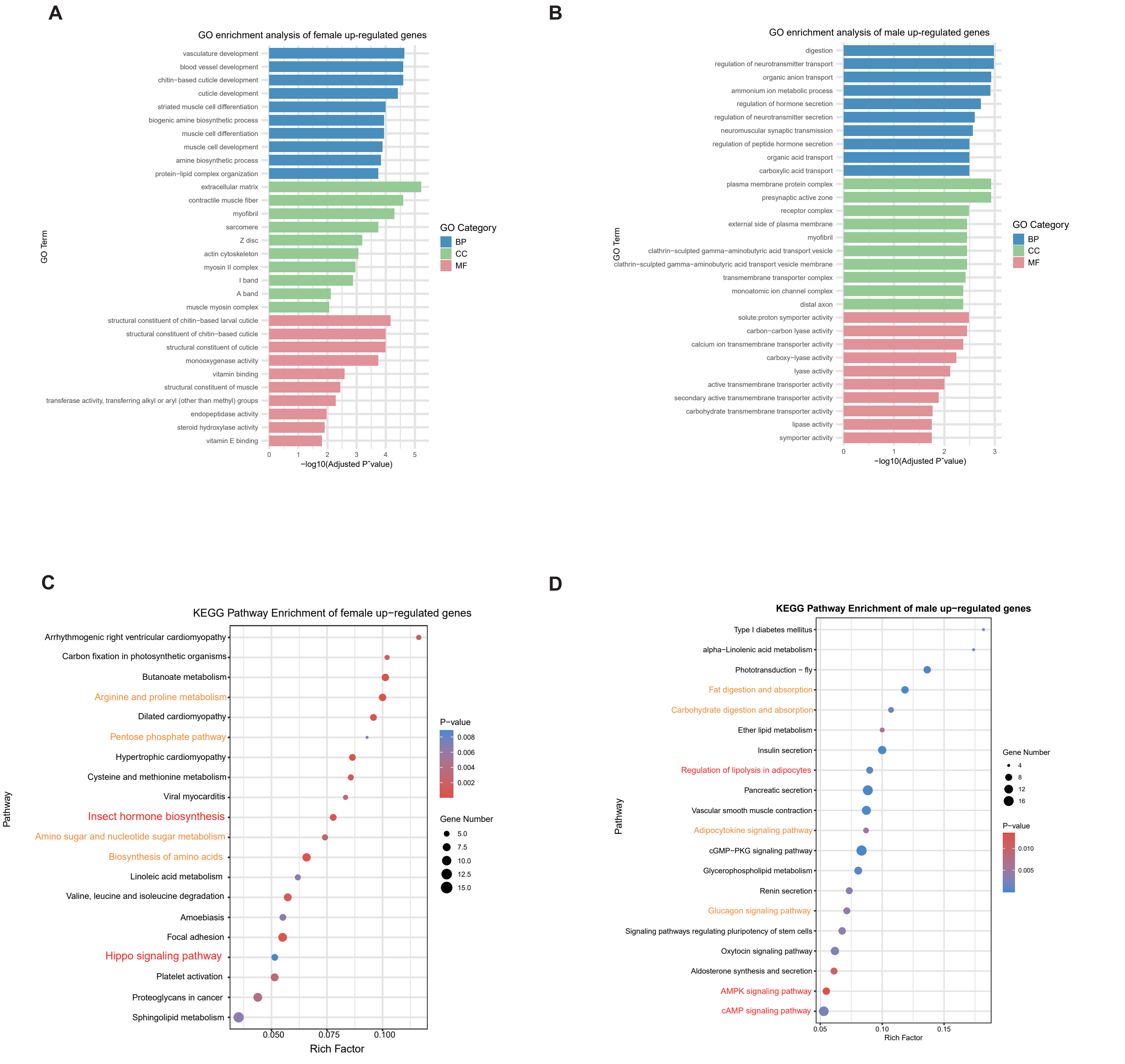


**Figure S2. Functional enrichment of sex-biased genes in adult *Trichonephila clavata*.**
(A − B) Gene Ontology (GO) enrichment of female-upregulated (A) and male-upregulated (B) genes. GO terms are grouped into biological process (BP), cellular component (CC), and molecular function (MF).(C − D) KEGG pathway enrichment of female-upregulated (C) and male-upregulated (D) genes. Dot size indicates gene number, and color indicates statistical significance.

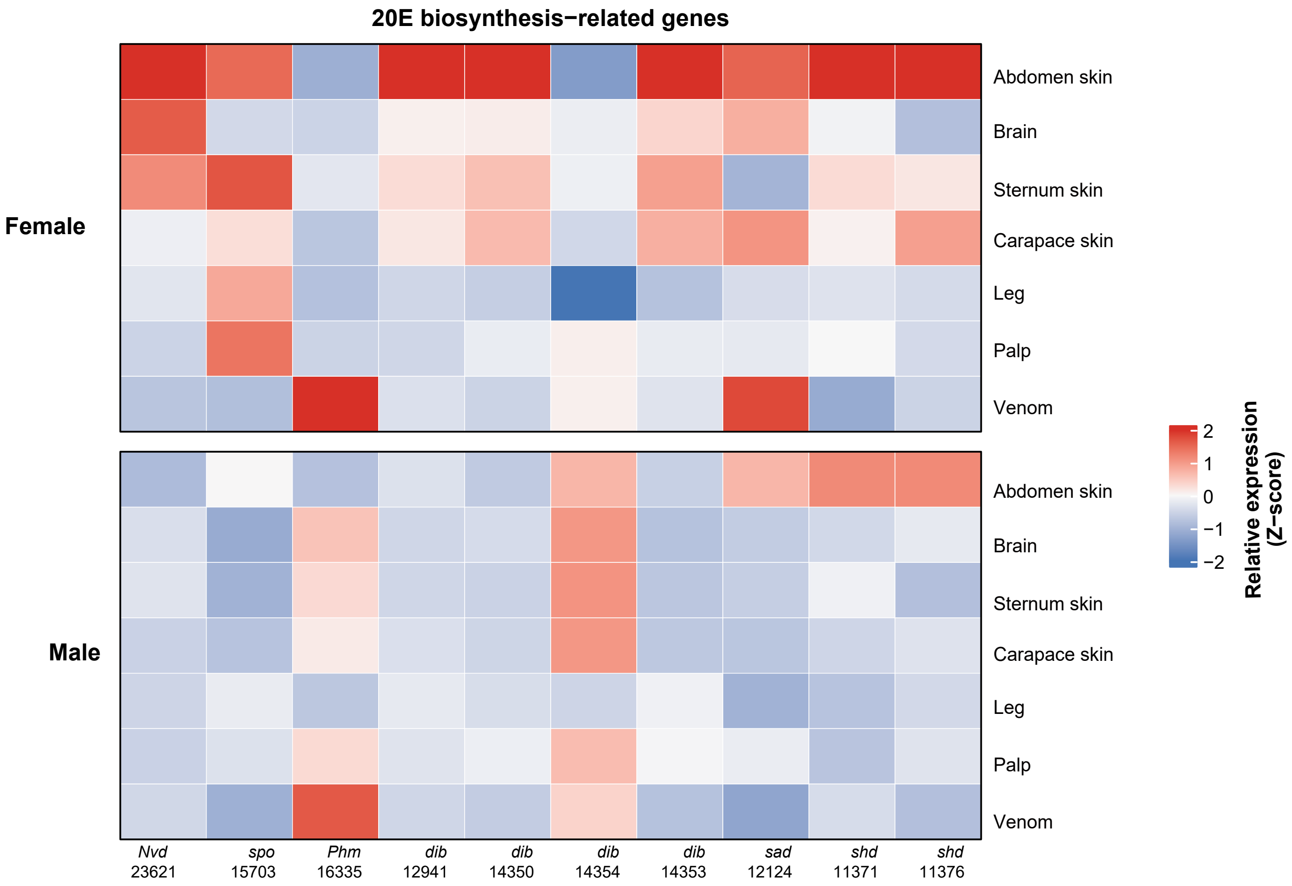


Figure S3. Tissue- and sex-dependent expression of 20E biosynthesis-related genes in adult *Trichonephila clavata*.

**Female**

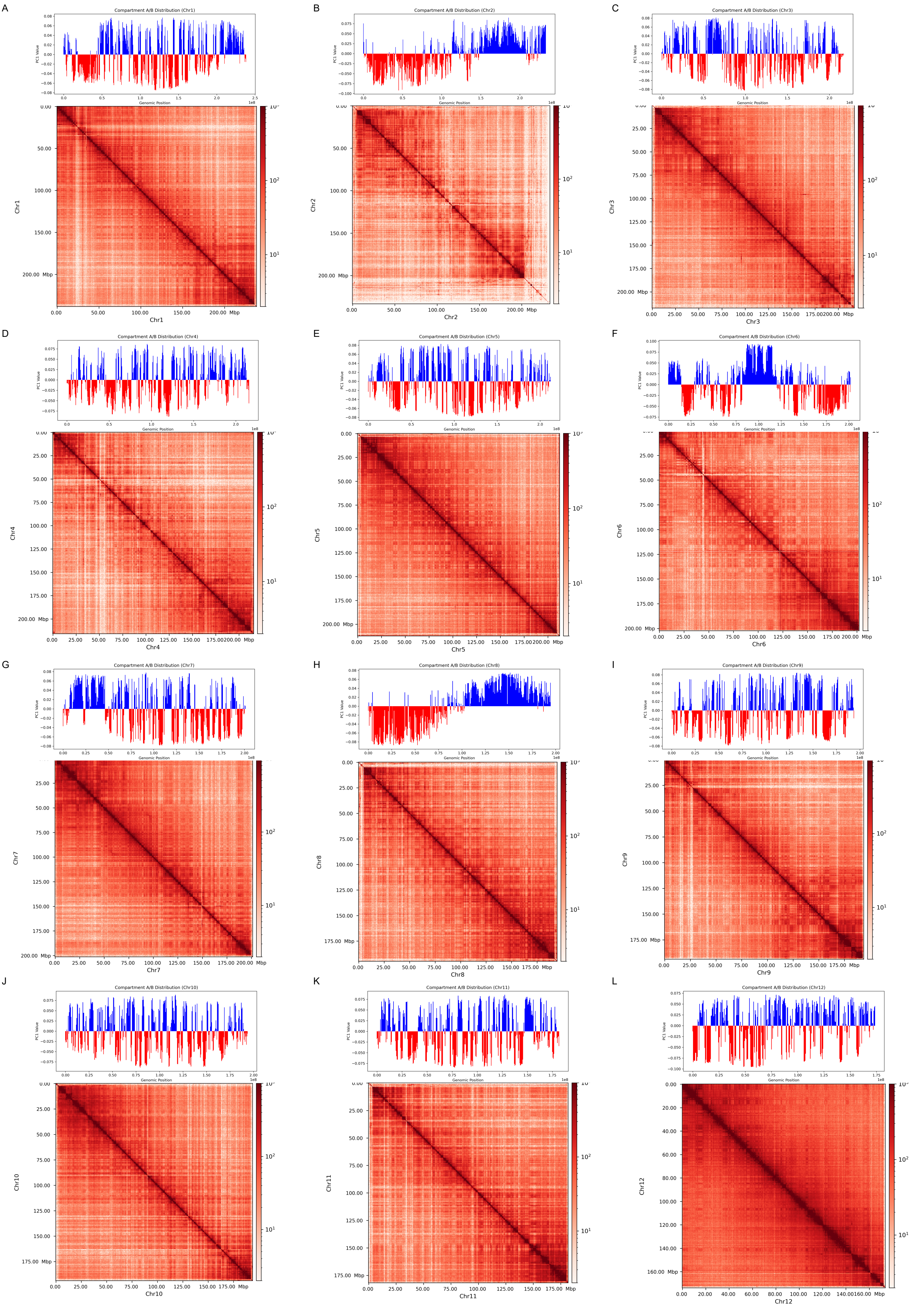
A
Compartment A/B Distribution (Chr1)
PC1 Value
Genomic Position
Chr1
B
Compartment A/B Distribution (Chr2)
PC1 Value
Genomic Position
Chr2
C
Compartment A/B Distribution (Chr3)
PC1 Value
Genomic Position
Chr3
D
Compartment A/B Distribution (Chr4)
PC1 Value
Genomic Position
Chr4
E
Compartment A/B Distribution (Chr5)
PC1 Value
Genomic Position
Chr5
F
Compartment A/B Distribution (Chr6)
PC1 Value
Genomic Position
Chr6
G
Compartment A/B Distribution (Chr7)
PC1 Value
Genomic Position
Chr7
H
Compartment A/B Distribution (Chr8)
PC1 Value
Genomic Position
Chr8
I
Compartment A/B Distribution (Chr9)
PC1 Value
Genomic Position
Chr9
J
Compartment A/B Distribution (Chr10)
PC1 Value
Genomic Position
Chr10
K
Compartment A/B Distribution (Chr11)
PC1 Value
Genomic Position
Chr11
L
Compartment A/B Distribution (Chr12)
PC1 Value
Genomic Position
Chr12
Mbp
1e8

**Figure S4. Chromosome-scale Hi-C interaction maps of female *Trichonephila clavata*.**
Genome-wide Hi-C contact matrices for chromosomes 1-12 in female individuals.

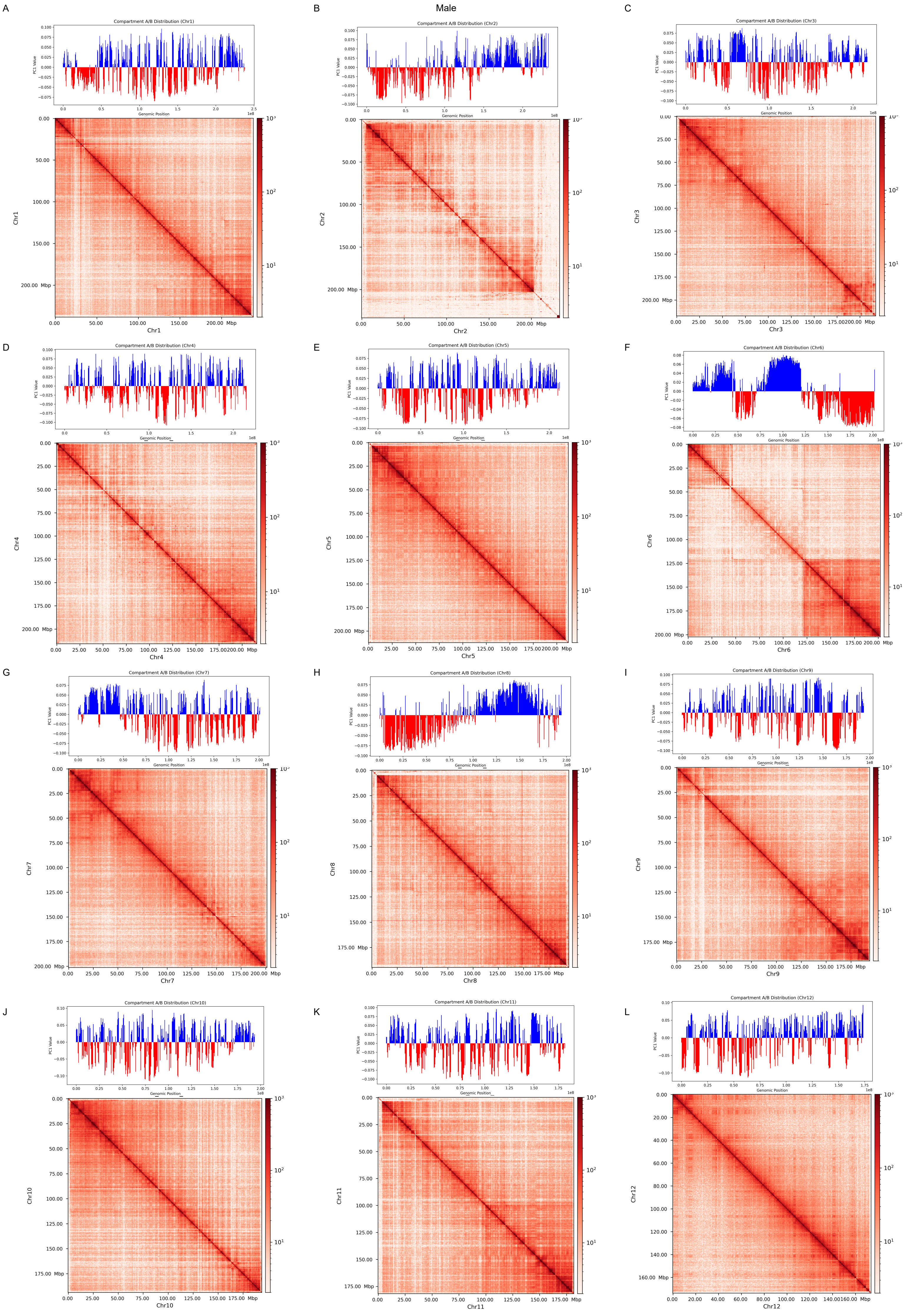
Male
A
Compartment A/B Distribution (Chr1)
B
Compartment A/B Distribution (Chr2)
C
Compartment A/B Distribution (Chr3)
D
Compartment A/B Distribution (Chr4)
E
Compartment A/B Distribution (Chr5)
F
Compartment A/B Distribution (Chr6)
G
Compartment A/B Distribution (Chr7)
H
Compartment A/B Distribution (Chr8)
I
Compartment A/B Distribution (Chr9)
J
Compartment A/B Distribution (Chr10)
K
Compartment A/B Distribution (Chr11)
L
Compartment A/B Distribution (Chr12)
PC1 Value
Genomic Position
1e8
Mbp
Chr1
Chr2
Chr3
Chr4
Chr5
Chr6
Chr7
Chr8
Chr9
Chr10
Chr11
Chr12

**Figure S5. Chromosome-scale Hi-C interaction maps of male *Trichonephila clavata*.**
Genome-wide Hi-C contact matrices for chromosomes 1-12 in male individuals.

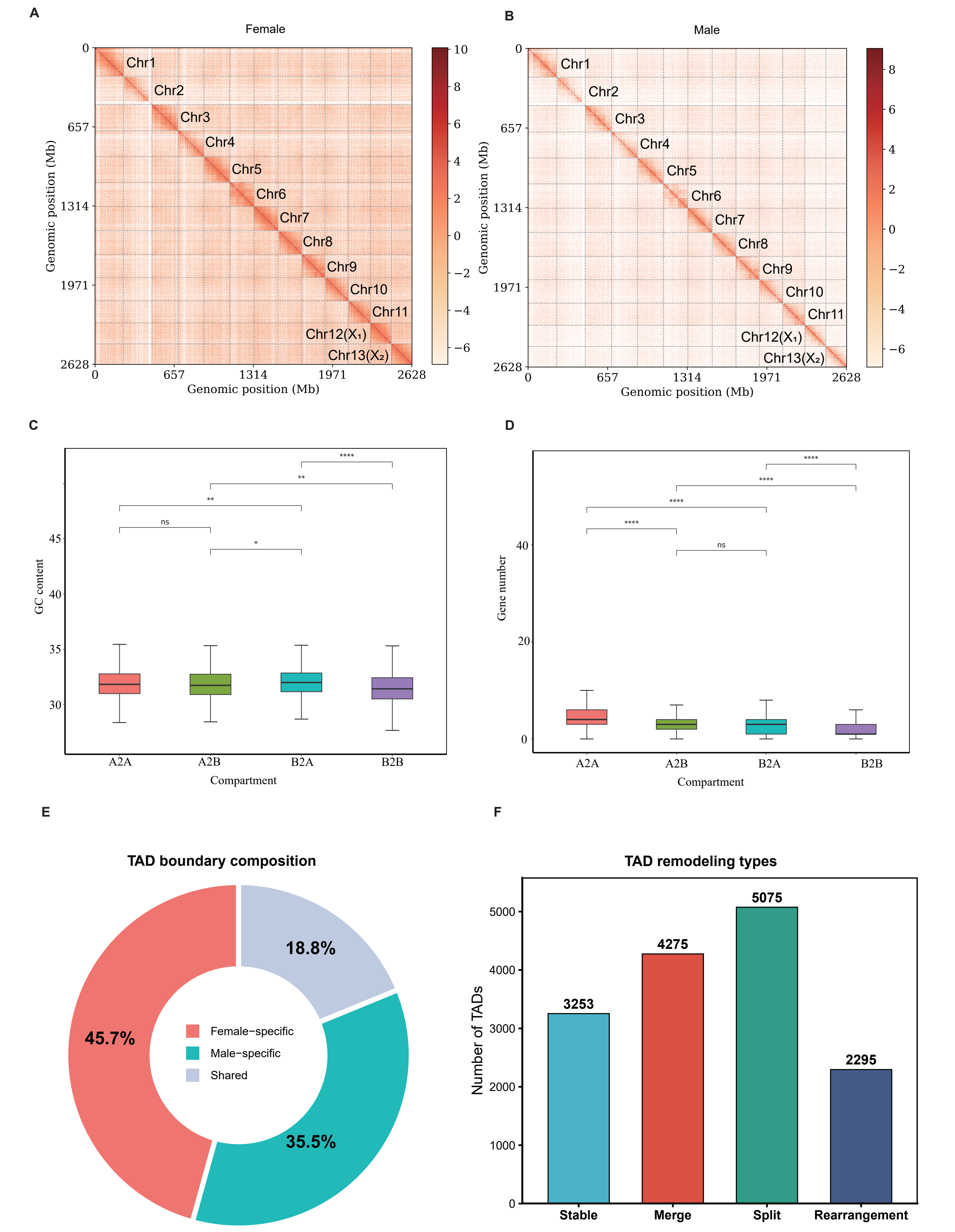


**Figure S6. Sex differences in three-dimensional genome organization in adult *Trichonephila clavata*.**
(A– B) Genome-wide Hi-C contact maps of females (A) and males (B).(C– D) GC content (C) and gene number (D) across regions classified by female-to-male A/B compartment states. Statistical significance is indicated above the comparisons.(E) Proportions of female-specific, male-specific, and shared TAD boundaries.(F) Numbers of TADs classified as stable, merged, split, or rearranged between females and males.

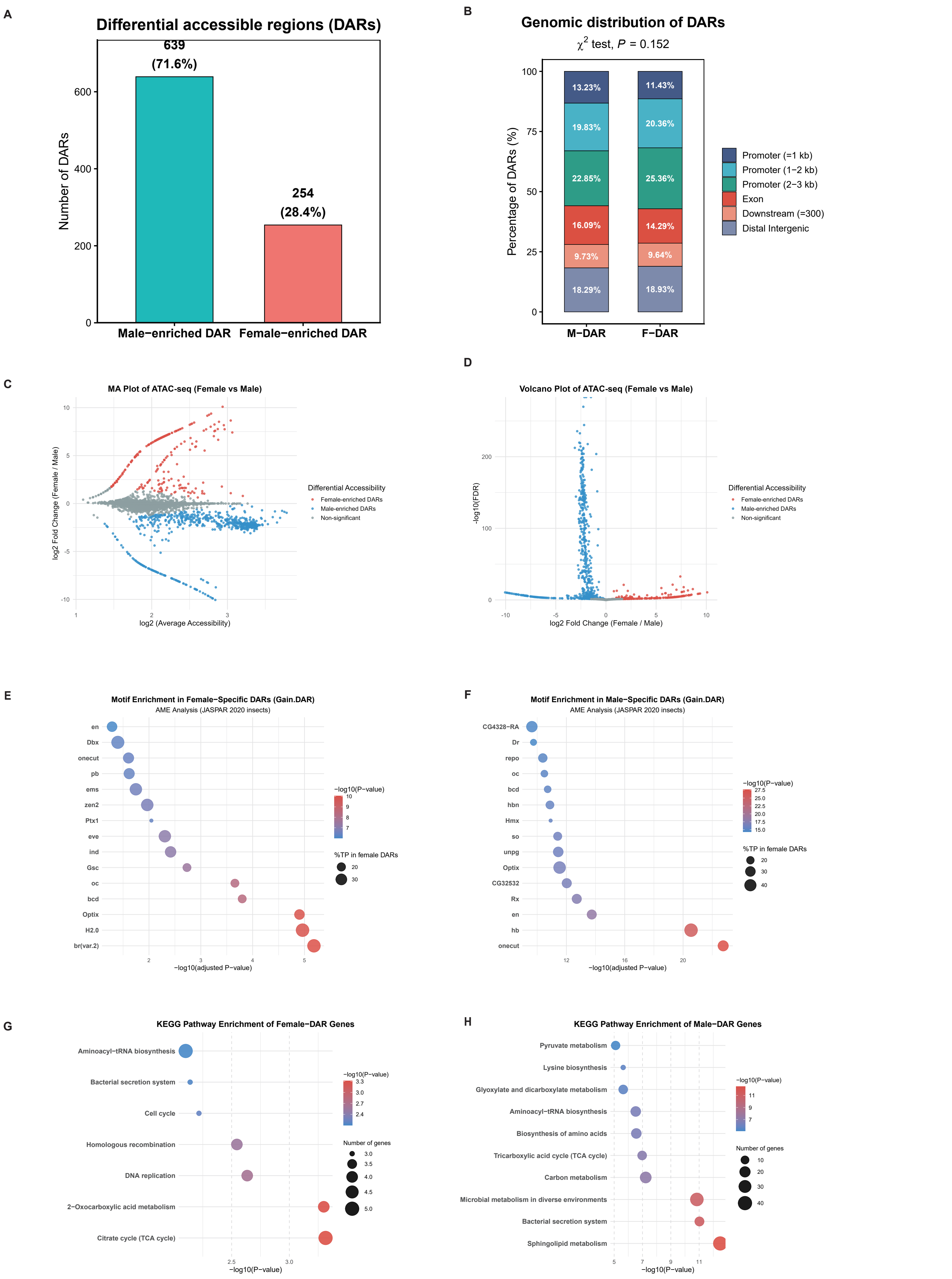


**Figure S7. Sex-biased chromatin accessibility landscapes revealed by ATAC-seq.**
(A) Numbers of differentially accessible regions (DARs) between females and males. (B) Genomic distribution of male-enriched (M-DARs) and female-enriched DARs (F-DARs) . (C) MA plot showing differential chromatin accessibility between females and males. (D) Volcano plot of differential chromatin accessibility between females and males. (E) Transcription factor motif enrichment analysis of female-specific DARs. (F) Transcription factor motif enrichment analysis of male-specific DARs. (G) KEGG pathway enrichment analysis of genes associated with female-enriched DARs. (H) KEGG pathway enrichment analysis of genes associated with male-enriched DARs.

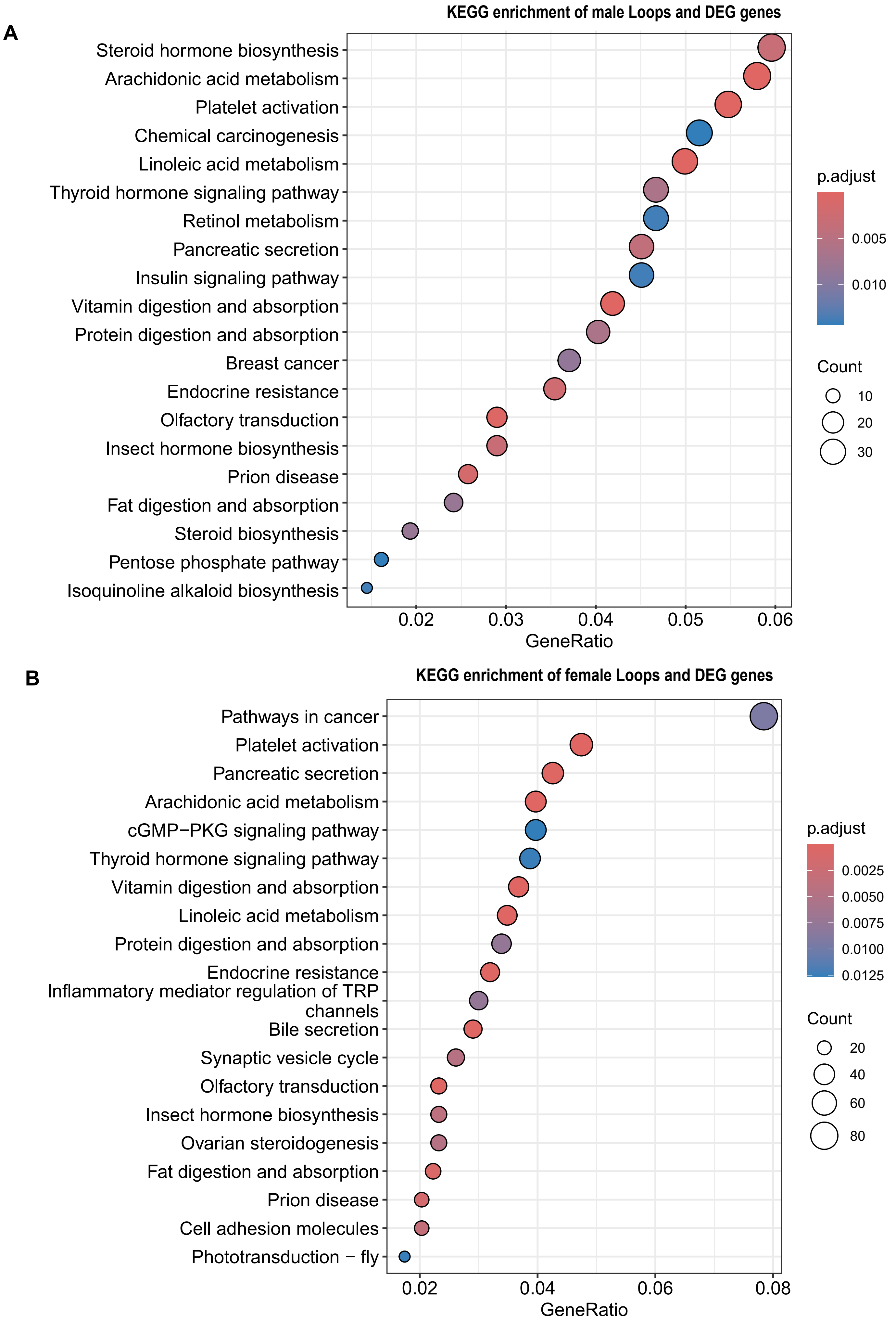


Figure S8. KEGG pathway enrichment analysis of male-biased genes (A) and female-biased (B) supported by both differential chromatin looping and differential gene expression (DEG).

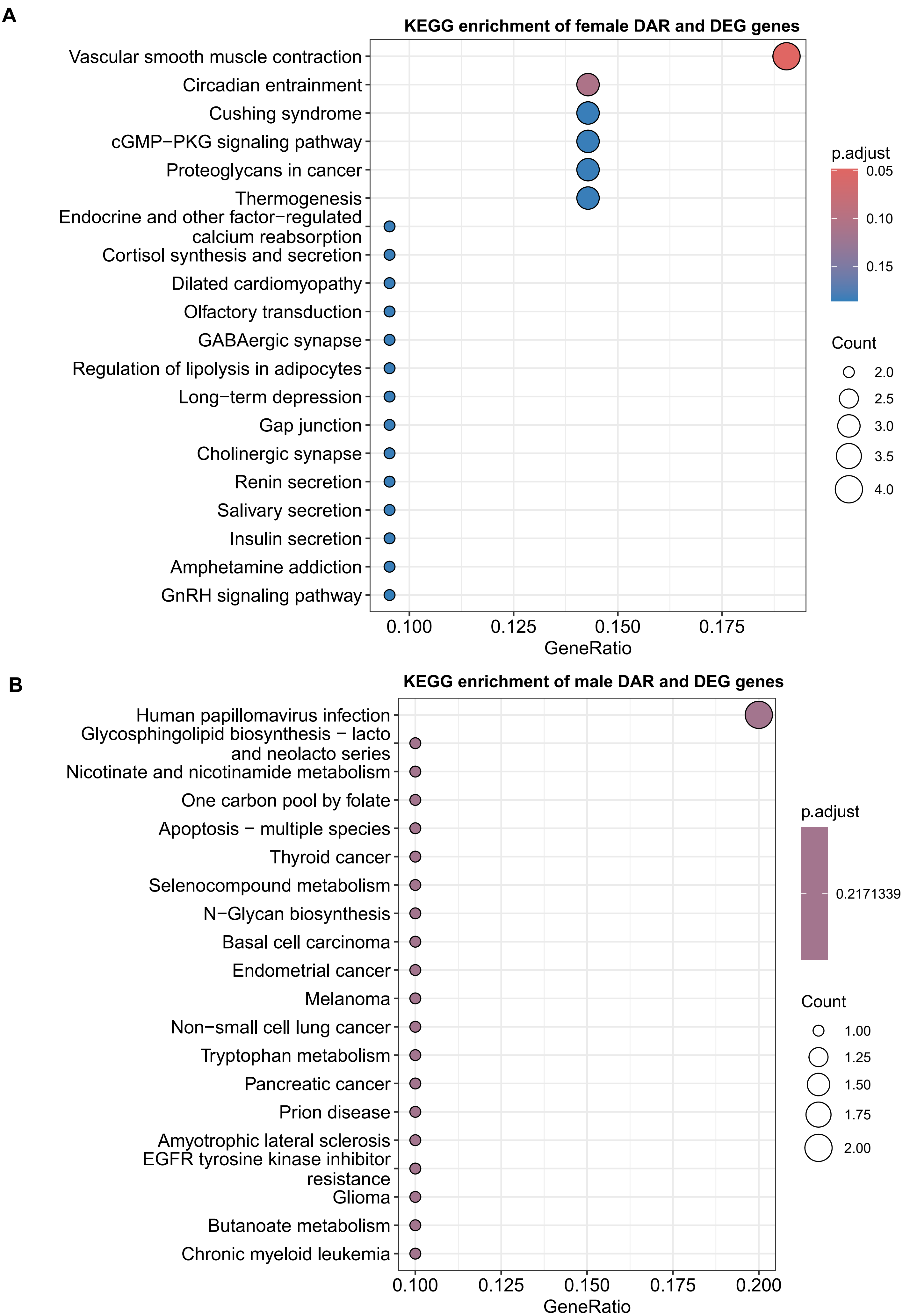


Figure S9. KEGG pathway enrichment analysis of male-biased genes (A) and female-biased (B) supported by both differential accessible regions (DARs) and differential gene expression (DEG).

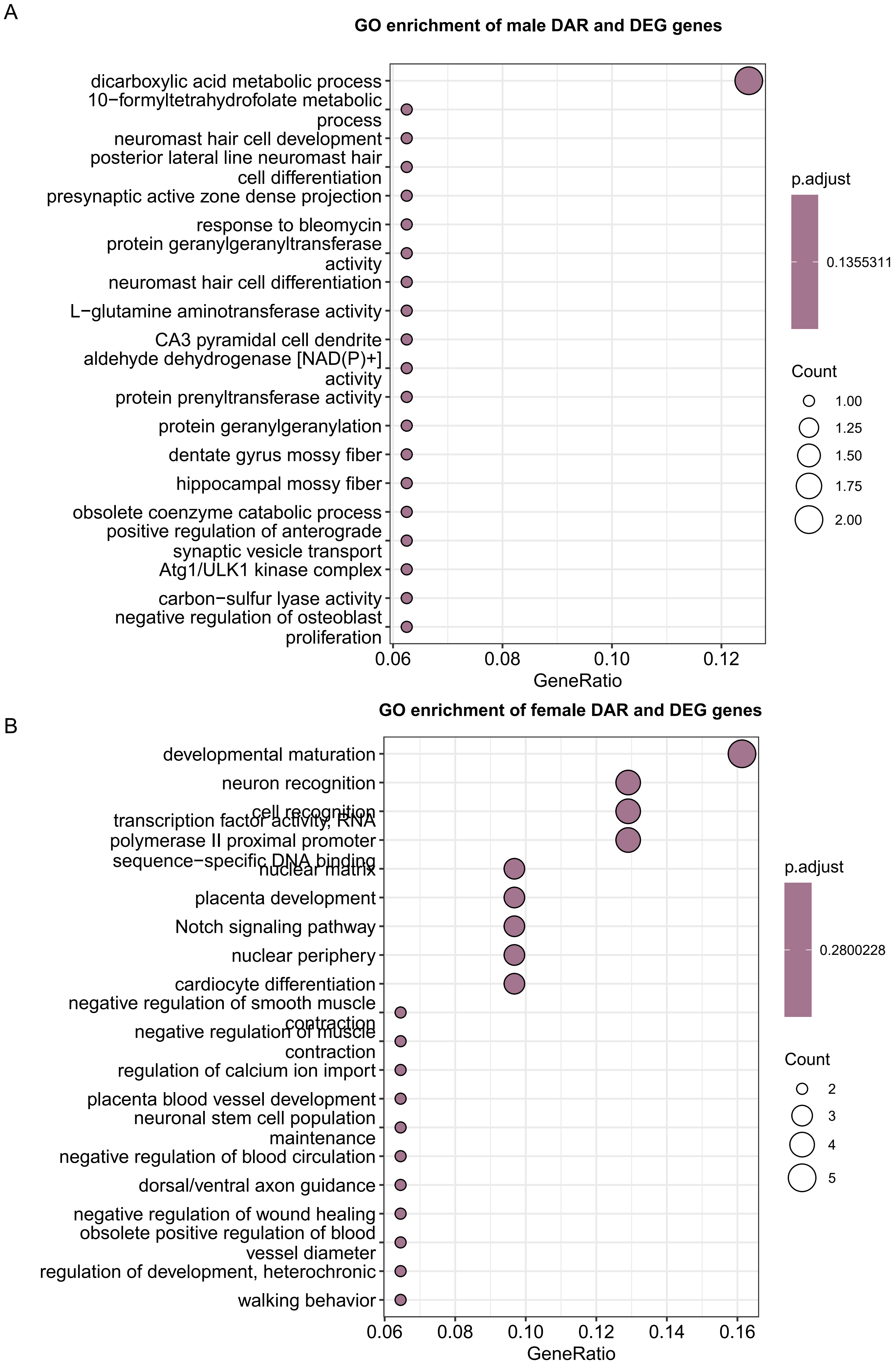


Figure S10. GO enrichment analysis of male-biased genes (A) and female-biased (B) supported by both differential accessible regions (DARs) and differential gene expression (DEG).

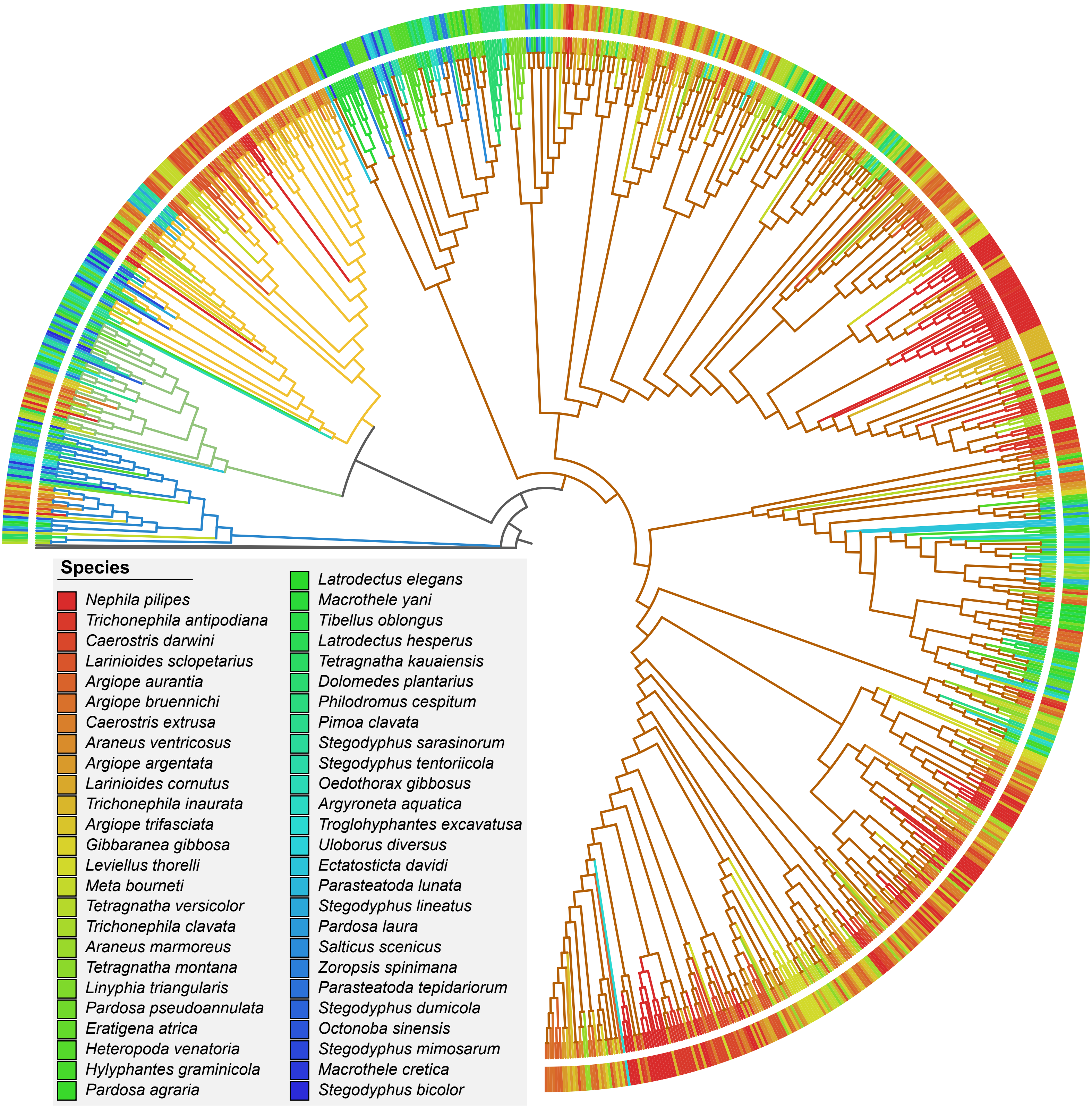


Figure S11. Tree of the juvenile hormone acid methyltransferase (JHAMT) gene family across 51 spider species.

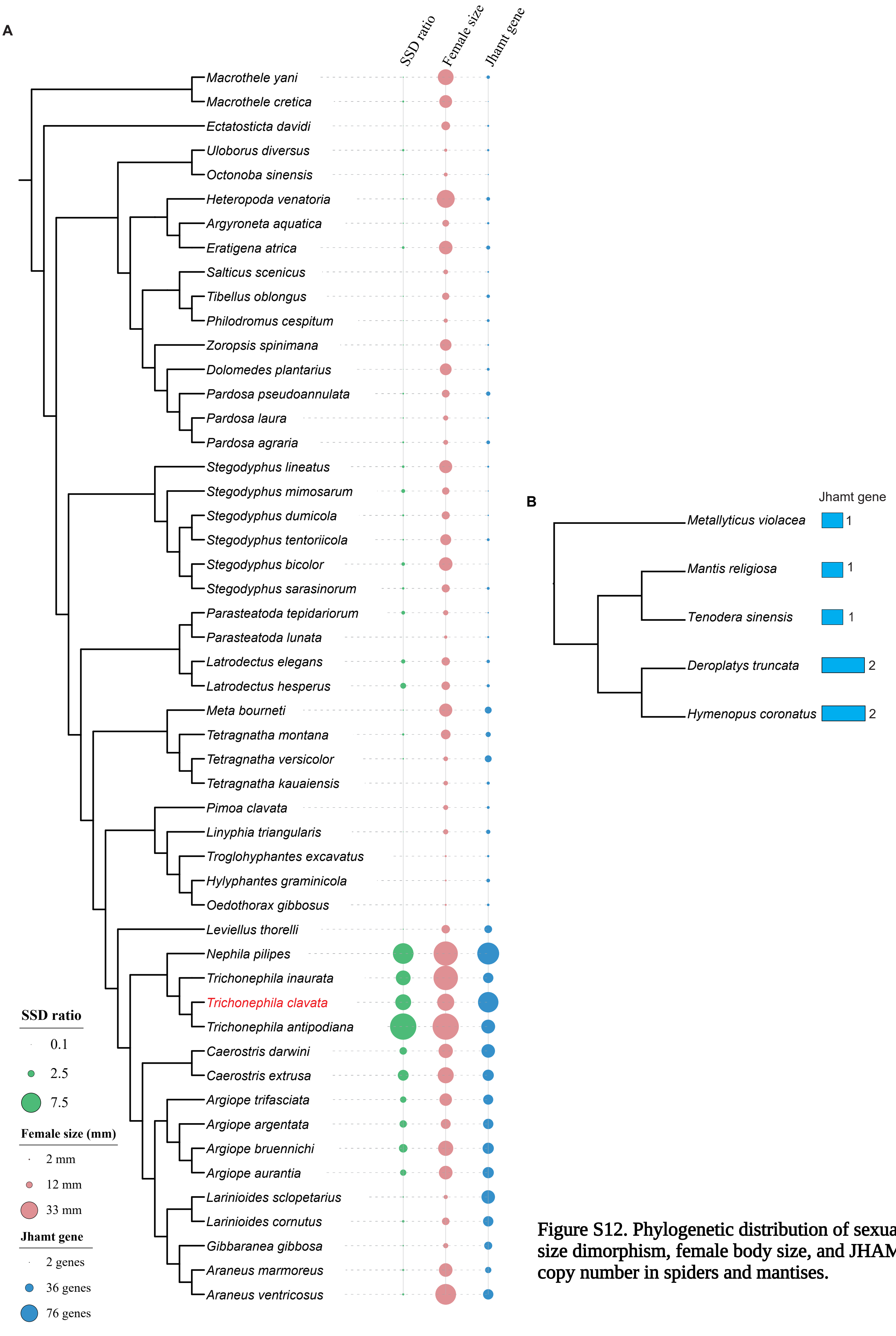


Figure S12. Phylogenetic distribution of sexual size dimorphism, female body size, and JHAMT copy number in spiders and mantises.

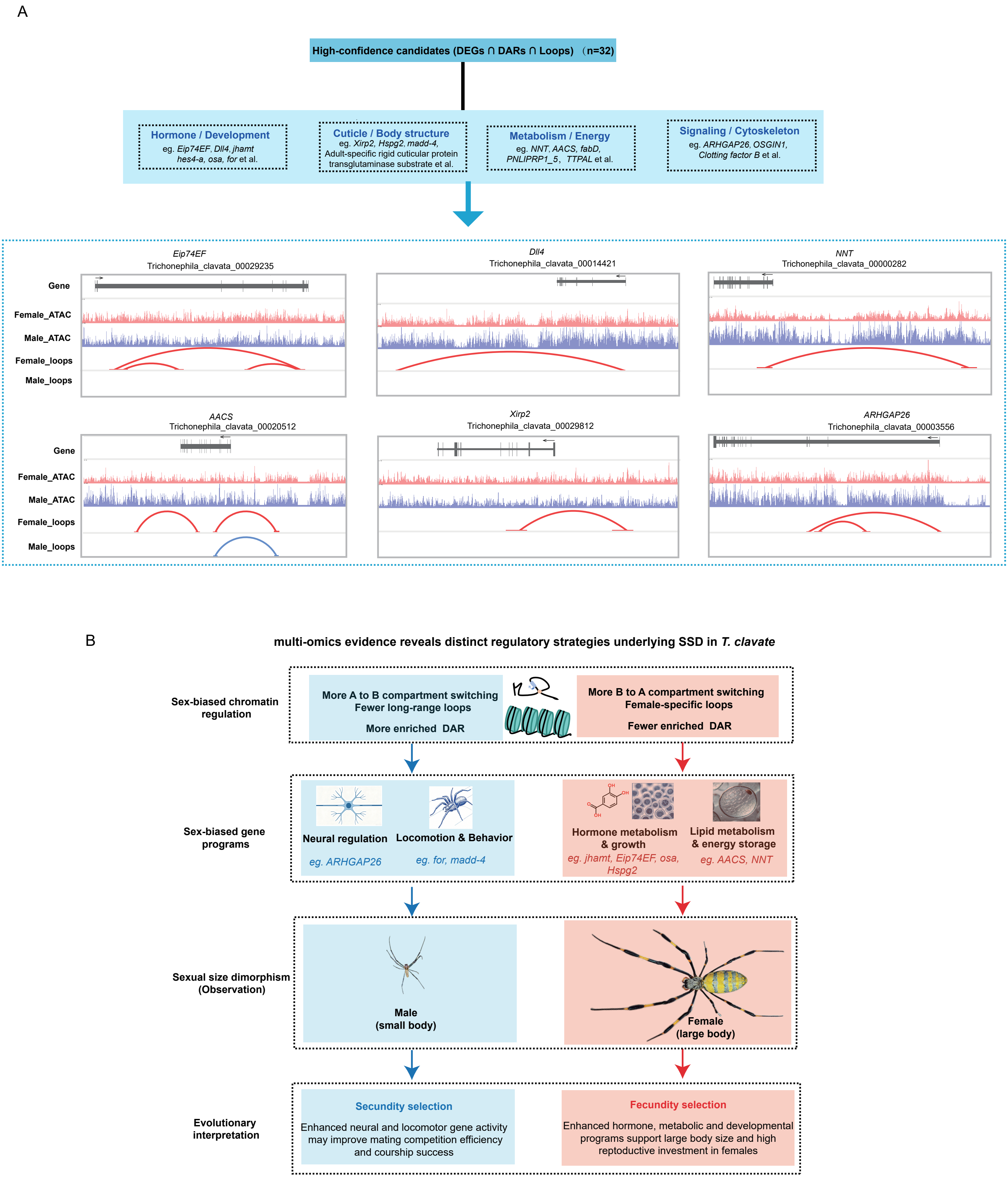


**Figure S13. Multi-omics integration identifies candidate regulatory genes associated with sexual size dimorphism in *Trichonephila clavata*.**

(A) Integration of differentially expressed genes (DEGs), differentially accessible regions (DARs), and loop-associated genes identified 32 high-confidence candidates.

(B) Working model summarizing sex-biased regulatory programs associated with SSD in *T. clavata*.
